# Including imprecisely georeferenced specimens improves accuracy of species distribution models and estimates of niche breadth

**DOI:** 10.1101/2021.06.10.447988

**Authors:** Adam B. Smith, Stephen J. Murphy, David Henderson, Kelley D. Erickson

**Affiliations:** Center for Conservation and Sustainable Development, Missouri Botanical Garden, 4344 Shaw Boulevard, Saint Louis MO 63110 USA; The Ohio State University, Department of Evolution, Ecology, and Organismal Biology, 318 W 12th Avenue, Columbus, OH 43210 USA; Department of Biology, Washington University in Saint Louis, 1 Bookings Drive, Saint Louis MO 63130 USA

**Keywords:** climate change vulnerability, coordinate uncertainty, georeferencing, niche breadth, natural history museum specimen records, niche truncation, niche estimation, rare species

## Abstract

**Aim:** Museum and herbarium specimen records are frequently used to assess species’ conservation status and responses to climate change. Typically, occurrences with imprecise geolocality information are discarded because they cannot be matched confidently to environmental conditions, and are thus expected to increase uncertainty in downstream analyses. However, using only precisely georeferenced records risks undersampling of species’ environmental and geographic distributions. We present two related methods to allow the use of imprecisely georeferenced occurrences in biogeographic analysis.

**Innovation:** Our two procedures assign imprecise records to the 1) locations or 2) climates that are closest to the geographic or environmental centroid of the precise records of a species. For virtual species, including imprecise records alongside precise records improved the accuracy of ecological niche models projected to the present and the future, especially for species with ^~^20 or fewer precise occurrences. Using only precise records underestimates loss in suitable habitat and overestimates the amount of suitable habitat in both the present and future. Including imprecise records also improves estimates of niche breadth and extent of occurrence. An analysis of 44 species of North American *Asclepias* (Apocynaceae) yielded similar results.

**Main conclusions:** Existing studies examining the effects of spatial imprecision compare outcomes based on precise records to the same records with spatial error added to them. However, in real-world cases, analysts possess a mix of precise and imprecise records and must decide whether to retain or discard the latter. Discarding imprecise records can undersample species’ geographic and environmental distributions and lead to mis-estimation of responses to past and future climate change. Our method, for which we provide a software implementation in the enmSdmX package for R, is simple to employ and can help leverage the large number of specimen records that are typically deemed “unusable” because of spatial imprecision in their geolocation.

## Introduction

Accurate estimation of species’ environmental tolerances and distributions is key to addressing many pressing issues in ecology, evolution, and conservation (e.g., Fisher-Reid et al. 2012; Quintero & Wiens 2013a and b; Foden & Young 2016). Information on species’ environmental tolerances and distributions is commonly inferred from occurrence records obtained from natural history museums and herbaria which are matched to associated environmental conditions. These conditions are then used to calculate the range of environments inhabited by a species (e.g., Foden et al. 2013). Similarly, ecological niche models (ENMs; also known as species distribution models) can be employed to estimate niche breadth and predict the extent of suitable habitat (Dawson et al. 2011; Miller et al. 2012; Pacifici et al. 2015; Foden & Young 2016; Young et al. 2016; IUCN 2019). Conservation assessments also frequently utilize the area encompassed by occurrence records to appraise species’ potential exposure to broad-scale threats (Faber-Langendoen et al. 2012; IUCN 2019). Methods utilizing opportunistic occurrence records have thus become an important component of biodiversity and conservation research (Heberling 2020). Unfortunately, a substantial portion of these records is imprecisely geolocated (Moudrý & Devillers 2020; Marcer et al. 2022), which poses challenges for estimating niches and distributions (Moudrý & Šímová 2012).

For occurrence data to be useful for estimating niche limits and distributions, two conditions are desirable. First, environmental information assigned to an occurrence record should reflect the conditions that were actually experienced by the organism at the location where it was observed (Graham et al. 2008). When records can only be geolocated imprecisely (i.e., to a large region or geopolitical unit), it becomes difficult to confidently assign a specific environmental datum to each record (Feeley & Silman 2010). Second, when the goal is to estimate environmental tolerances, the set of records used to estimate niche limits should encompass as much of the species’ fundamental niche as possible (Thuiller et al. 2004; Peterson et al. 2018). If available occurrences represent only a portion of the niche, then the species’ actual environmental tolerances will be underestimated (Thuiller et al. 2004; Qiao et al. 2017).

The desire for spatial precision and representative sampling of occurrences leads to a critical trade-off. On the one hand, if records can only be imprecisely geolocated, using them risks introducing uncertainty into estimates of environmental tolerances (Graham et al. 2008; Fernandez et al. 2009; Osborne & Leitão 2009; Feeley & Silman 2010; Tulowiecki et al. 2015; Gábor et al. 2020 and 2022; Mitchell et al. 2016; Collins et al. 2017; Cheng et al. 2021; Marcer et al. 2022). On the other hand, discarding records risks under-representing the true geographic and environmental range of a species, even if the location of occurrences is uncertain (Graham et al. 2008). These risks are especially great for rare species, which tend to be represented by just a few records (Lomba et al. 2010; Sheth et al. 2012; Zizka et al. 2018).

To date, these trade-offs have been almost exclusively managed in favor of reducing apparent uncertainty by discarding imprecisely georeferenced records (Moudrý & Šímová 2012). Indeed, in a literature survey of peer-reviewed publications using museum or herbarium specimens with ENMs, we found that of studies that described their data cleaning process, 45% discarded spatially imprecise records (Fig. 1; Appendix S1). No studies reported purposefully retaining them or using methods to accommodate imprecision. In other words, the risk of including imprecise occurrence data has been assumed more serious than undersampling of the realized niche. However, the justification for discarding spatially imprecise records is often based on studies that do not necessarily reflect real-world use cases. Instead, these studies evaluate the effects of coordinate imprecision on the accuracy of ENMs by artificially adding spatial error to otherwise precise records, and then comparing results between “fuzzed” and accurate records (e.g., Graham et al. 2008; Fernandez et al. 2009; Osborne & Leitão 2009; Gueta & Carmel 2016; Mitchell et al. 2016; Hefley et al. 2017; Soultan & Safi 2017; Tulowiecki et al. 2015; Gábor et al. 2020 and 2022). This approach, however, is not reflective of the common situation where an assessor starts with a mix of relatively precisely- and imprecisely-geolocated records and must decide how to delineate the two groups and whether or not to retain the imprecise ones (cf. Marcer et a. 2022).

**Figure 1.**
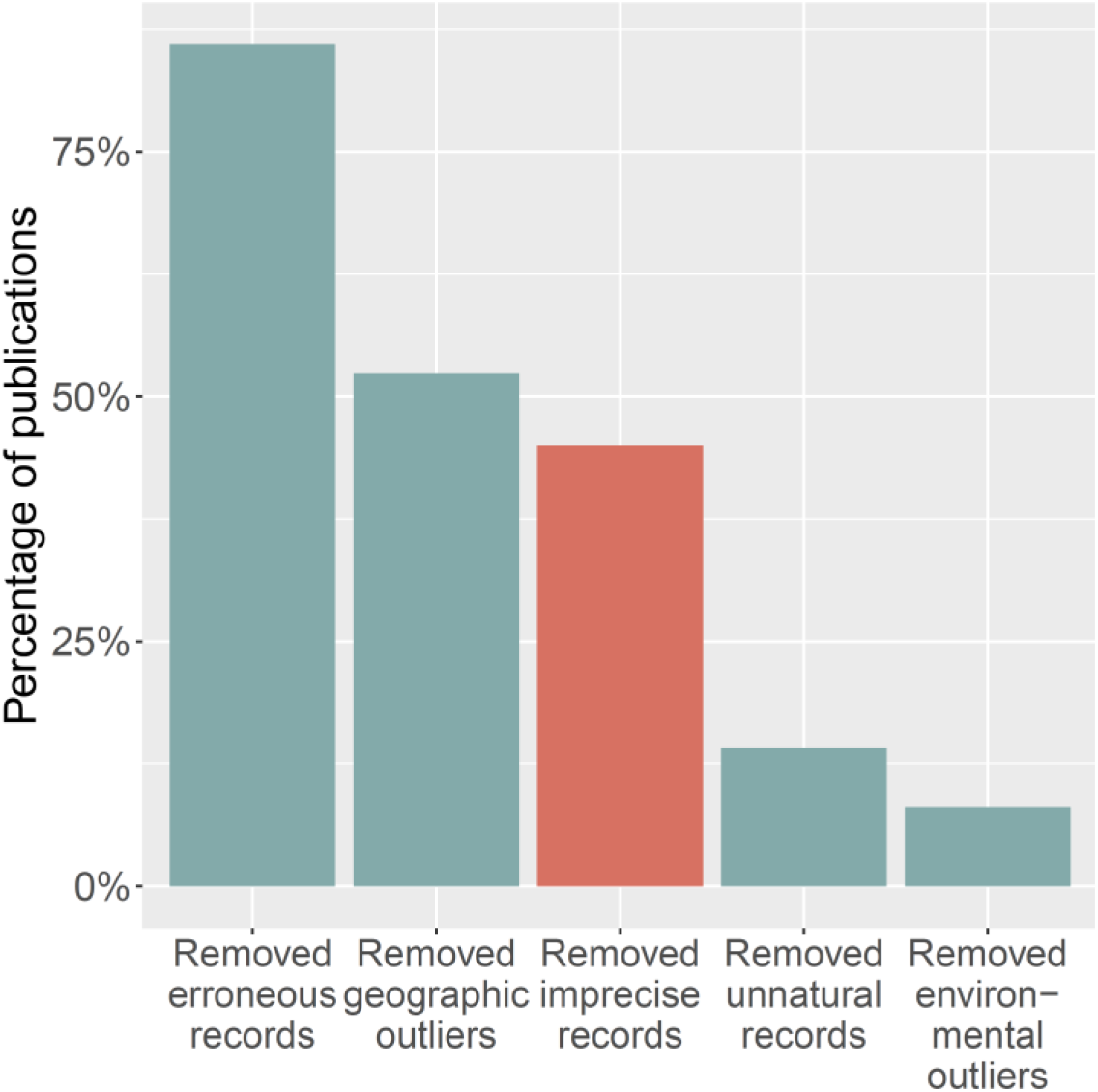
Results of literature survey on how imprecise specimen records are treated in peer-reviewed publications that used herbarium or museum specimens, employed species distribution models, and were published between 2010 and 2019. Of 285 relevant, randomly selected studies, only 149 (52%) described any data cleaning procedures. Bars represent the frequency of studies that report the given data cleaning method, as a percentage of studies that described any data cleaning procedure (see Appendix S1 for description of categories, and sampling and scoring methods).

Here we present two simple, related methods for incorporating imprecise records into biogeographic analyses. We use these methods to reexamine the trade-offs between retaining versus discarding spatially imprecise records using virtual and real species. Records were designated as “precise” if they had spatial uncertainty small enough to match them with confidence to environmental data, and “imprecise” if their spatial uncertainty was too great for confident assignment to a single environmental raster cell. Imprecision in record coordinates can be represented in a variety of ways, including through the use of user-defined polygons (Wieczorek et al. 2004) or by assignment of records to the smallest geopolitical unit encompassing the area of likely collection (e.g., county, province, etc.; Park & Davis 2017). We emphasize that our definition of an imprecise record does not include records with locations appearing to be outside the range of the species (i.e., geographic outliers; Feeley & Silman 2010) or specimens that do not pass quality-assurance checks (Chapman 2005). We compared ENMs, niche breadth, and the spatial extents of occurrence estimated using only precise records to estimates based on precise plus imprecise records. We evaluated the accuracy of each of these metrics as a function of how many imprecise records were included, and compared them to benchmark estimates based on “omniscient” records where all populations of a species could be georeferenced without error.

## Methods

Imprecise records can, by definition, be associated with multiple locales. This poses a challenge, since the choice of a specific locale will affect the apparent spatial and environmental distribution a species. Implemented in geographic space, we call our solution to this problem the “nearest geographic point” (NGP) method (Fig. 2a). We believe the most useful application of NGP is calculation of extent of occurrence (EOO), the area of the minimum convex polygon circumscribing all known populations. EOO is commonly used in conservation assessments to assess exposure to widely-distributed threats (Young et al. 2016; IUCN 2019). To implement NGP we start by 1) locating the geographic centroid of precise records. Imprecise records are assumed to be represented by spatial polygons of the area of likely collection (e.g., an administrative unit). So, we then 2) find for each such polygon the point on its border that is closest to the centroid from step 1. To construct the minimum convex polygon for estimating EOO, we then 3) use the combined set of precise records plus the points identified in step 2 from the imprecise records. The actual location of collection an imprecise record will almost always be further from the precise records’ centroid than the closest location on an imprecise record’s polygon. Hence, this method will tend to yield conservative estimates of EOO compared to, say, using the centroid of polygons representing areas of likely collection (e.g., Collins et al. 2017; Park & Davis 2017).

**Figure 2.**
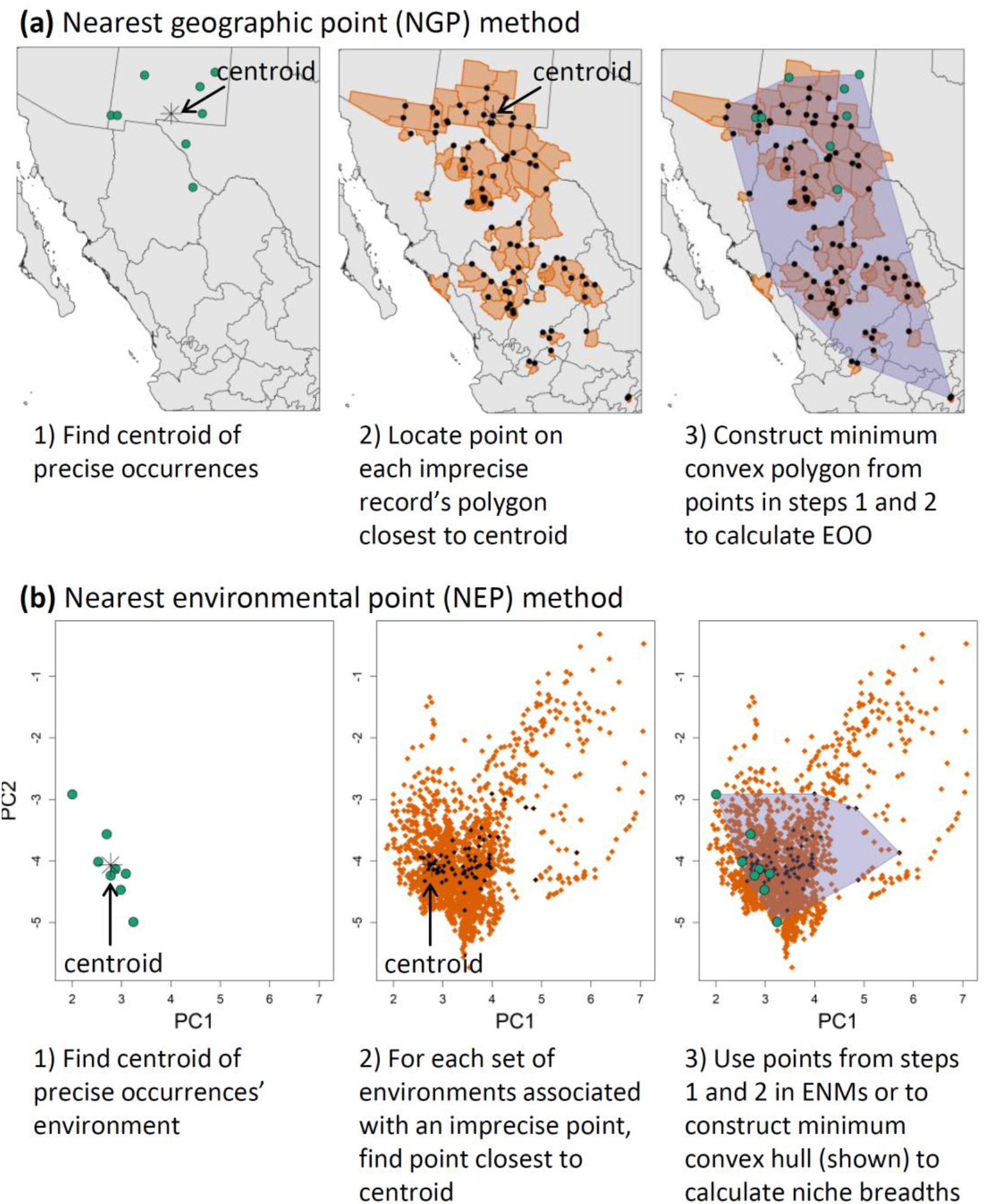
Illustrations of how to apply the (a) “nearest geographic point” and (b) “nearest environmental point” methods for incorporating imprecise records into biogeographic analyses. Analysis is shown using data for *Asclepias brachystephana*. In each case, green circles represent precise occurrences. Orange polygons (a) or points (b) represent county-level occurrences or the set of environments associated with these counties, respectively. Calculations for NEP are shown for a 2-dimensional space defined by principal component axes, but NEP can be applied in any number of dimensions.

For analyses in environmental space, we use a related procedure, the “nearest environmental point” (NEP) method. NEP is similar to NGP, except that it can be implemented in one or more dimensions (one or more environmental axes). NEP starts with 1) finding the mean value of environmental conditions across precise records (Fig. 2b). Since imprecise records can only be located to a general area (e.g., a county), for each imprecise record we then 2) extract all environments across this area, then find among them the one that is closest to the mean value from step 1. These environmental values, along with the values from the precise records, are 3) then used in downstream analyses (e.g., for calculating niche breadth or calibrating ENMs). If the environmental space is multi-dimensional and variables are in different units, we first transform them using a principal component analysis (PCA). Importantly, the environments assigned to imprecise records using NEP will not necessarily be the ones associated with the locations assigned to them using NGP because the environments identified with NEP will not necessarily be the ones geographically closest to the geographic centroid of the precise records. Hence, we do not use geographic locations identified using NGP as the locales from which environmental values are to be extracted. If analyses are to be conducted in both geographic and environmental space, NGP and NEP should be implemented separately.

To implement these methods, we developed the functions `mcpFromPointsPolys` and `nearestEnvs` available in the `enmSdmX` package (Smith 2022; https://cran.r-project.org/package=enmSdmX) for R (R Core Team 2021). Both functions can handle spatial objects from the terra (Hijmans 2022) and sf (Pebesma 2018) packages.

### Case studies using virtual and real species

For both case studies, the region of analysis encompassed North America (i.e., Canada, the United States, and Mexico, excluding distant islands such as the Hawaiian Islands). However, we calibrated and evaluated ENMs within species-specific regions as noted below.

#### Virtual species

We evaluated our methods using virtual species for which we know true distribution and environmental tolerances (Meynard et al. 2019). Fig. 3 illustrates the workflow. To create species, we first calculated a PCA on all present-day climatic conditions for North America using all 19 BIOCLIM variables (Nix 1986) from WorldClim Version 2.1 at 10 arcmin resolution, which represents average conditions across 1970-2000 (Fick & Hijmans 2017).

**Figure 3.**
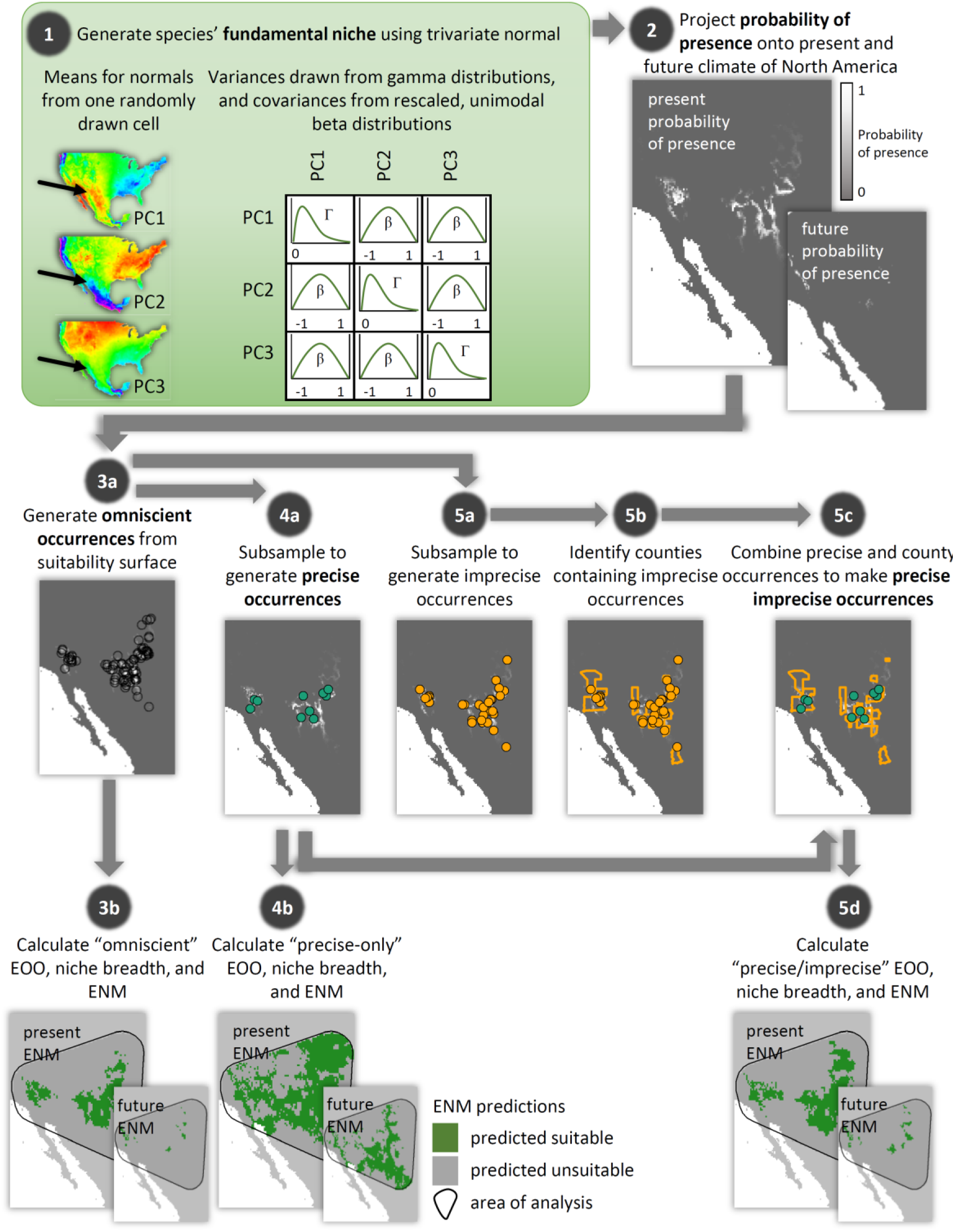
The process of generating and analyzing a virtual species starting with generation of the fundamental niche (step 1), projection to geographic space (2), and generation of omniscient (3), precise (4), and imprecise (5) occurrences. Suitable climate area is analyzed within a buffered region surrounding omniscient records (bottom row). The process was repeated 200 times for each combination of number of omniscient, precise, and imprecise records.

We defined the fundamental niche of each species as a multivariate normal distribution using the dmvnorm function in the mvtnorm package (Genz & Bretz 2009; Genz et al. 2021). To parameterize the fundamental niche for a species, we drew at random a cell with a probability proportionate to the cell area from across Mexico and the US, then used the scores from the first three PC axes to specify the means for the multivariate normal function. We excluded Canada in this step because including it frequently resulted in a species that had few/no administrative records, since much of Canada is not divided into sub-provincial units. We drew a standard deviation for each variable from a gamma distribution with “shape” parameter of 1 and “rate” parameter itself drawn from a uniform distribution across (7.5, 30). We allowed for pairwise interactions between each variable by defining a covariance for each pair of variables drawn from a symmetrical, unimodal beta distribution with parameters α = β = 5, which tended to result in weak-to-moderate interactions between niche variables. Preliminary analyses revealed that these methods and parameterizations were capable of simulating a wide variety of species with ranges commensurate in extent to real species we analyzed (described below). We then generated a climatic suitability raster by projecting the multivariate normal function to the stack of rasters representing the three PC axes across North America.

We created a set of “omniscient” records, which represented all known and potentially unknown populations of a species. Omniscient records were located by drawing, without replacement, cells with a probability proportionate to their climatic suitability times each cell’s area. For each species, we generated 20, 40, 80, 160, or 320 omniscient records. Analyses using these records serve as a benchmark against which to compare analyses using precise and imprecise records.

To create “precise” records, we randomly sampled without replacement a subset of the omniscient records. These represented occurrences that collectors have georeferenced with minimal error. For each number of omniscient records, we selected a number of precise records along the series {5, 10, 15, 20, 25, and 30}, so long as *N_P_* < *N_O_* (where *N_P_* and *N_O_* represent the number of precise and omniscient records, respectively).

To create “imprecise” records, we sampled from the remainder of omniscient records. These were then matched to the county (or equivalent geopolitical unit) in which they were located. For each number of precise records, we selected a number of imprecise records along the doubling sequence {1, 2, 4, 8,…, up to *N_O_* – *N_P_*}. We generated 27 combinations of scenarios consisting of 5 levels of omniscient records (from 20 to 320 records), four of which were paired with 6 levels of precise records (from 5 to 30 records), except for the case where *N_O_* = 20, which had three levels (from 5 to 15 records). For each scenario, we generated 200 species.

For each species, we conducted an: a) “omniscient analysis” using all omniscient records; b) a “precise-only” analysis using just precise records; and c) a “precise+imprecise” analysis using the combined set of precise and imprecise records. For each analysis, we calculated several metrics relevant to conservation in geographic and environmental space. For the geographic analysis, we calculated EOO from the minimum convex polygon encompassing all records using the exact location of the records (omniscient and precise-only analysis) or using the precise records plus locations for imprecise records identified using NGP (precise+imprecise analysis).

For the environmental analysis, we compared estimates of niche breadth and exposure to future climate change from ENMs. For the omniscient and precise-only analyses, we extracted environmental variables from the exact locations of the occurrences. For the precise+imprecise analysis, we extracted environmental values from the exact locations of the precise records and combined them with environmental values extracted using NEP. We then calculated univariate niche breadth in mean annual temperature and total annual precipitation, using the range of each variable encompassed by the respective set of occurrences. We also estimated multivariate niche volume and surface area from the convex hull of the species’ occurrences in environmental space using the convhulln function in the geometry package (Habel et al. 2022).

We constructed ENMs using MaxEnt 3.4.3 (Phillips et al. 2006; Phillips & Dudík 2008) using the trainMaxEnt function in the enmSdmX package (Smith 2022). This function identifies the optimal combination of terms and master regularization parameter using AIC_c_ to reduce over/underfitting (Warren & Siefert 2011). We evaluated all possible combinations of linear, quadratic, and interaction terms plus values of the regularization parameter across{0.5, 1, 1.5, 2, 2.5, 3, 4, and 5}. Background sites for calibration of all models were drawn from a 300-km buffer around the minimum convex polygon constructed around all omniscient points. Models were projected to the present and to a future climate scenario defined by the average across five earth system models for 2061-2080 from CMIP6 under Representative Concentration Pathway 8.5 (achievable under Shared Socioeconomic Pathway 5-8.5; O’Neill et al. 2015).

We assessed calibration accuracy (the degree to which model output reflects the true probability of presence) of the model predictions in the present and the future using the Pearson correlation coefficient between the ENM predictions and the actual probability of presence. We also calculated the area of current and future favorable habitat, and the area of habitat that was lost, gained, or remained favorable after applying a threshold to the prediction maps such that training sensitivity was 0.9, then. Each of these assessments was conducted using either the area delineated by a 300-km buffer around the minimum convex polygon circumscribing all omniscient records, or 10000 random sites from this region. We measured model complexity as the number of non-zero coefficients.

We did not compare results between data types (omniscient, precise-only, precise+imprecise) using statistical hypothesis testing since these are inappropriate for simulations where the existence of differences is known a priori (White et al. 2014). Rather, we compared the inner 90^th^-percentile distribution of each metric (EOO, calibration accuracy, etc.) for each case.

#### Real species

We also evaluated 44 species of *Asclepias* (milkweeds; family Apocynaceae) native to North America, which display a range distributions from narrowly endemic to those covering approximately one-third of the continent. Records were obtained from the Global Biodiversity Information Facility (www.gbif.org), and detailed procedures for data cleaning and modeling are described in Appendices S3 and S4 (Zurell et al. 2020). We used only herbarium specimens and species with ≥5 precise, non-duplicate records. We evaluated the same set of metrics as for the virtual species and created ENMs following the same procedures. Since we did not have omniscient records for the real species, we estimated climatically suitable habitat area within the region defined by a 300-km buffer around the minimum convex polygon surrounding all available records. We tested for differences in EOO and in univariate and multivariate niche breadth calculated with or without imprecise records using a paired Wilcoxon signed-rank test.

#### Reproducibility

Scripts for the analyses are available on the GitHub repository https://github.com/adamlilith/enms_impreciseRecords. The analyses relied primarily on the `sp` (Bivand et al. 2013), `rgeos` (Bivand & Rundel 2020), geosphere (Hijmans 2019), `dismo` (Hijmans et al. 2022), raster (Hijmans 2022), and `enmSdmX` (Smith 2022) packages for R Version 4.10 (R Core Team 2021).

## Results

### Virtual species

We found that adding imprecise records caused the greatest improvements in metrics when the number of precise records was <15-20. So, for brevity, in the main text we focus on results for species with 40 and 320 omniscient records, each with 5 or 20 precise records (see Appendix S2 for all results). To reflect real-world use cases where assessors have a set of imprecise records but not otherwise know if they represent truly geographically unique specimens (e.g., different places in the same county), we plotted the change in each metric (ENM accuracy, niche breadth, etc.) against the number of county-level records accrued as imprecise records were added. Counties were only counted once, regardless of the number of imprecise records they contain. We thus hereafter describe trends in terms of number of county-level records added.

#### Extent of occurrence

Using only precise records consistently underestimated EOO (Fig. 4a and Appendix S2 Fig. 2.12). Underestimation was worst for species with few precise records and a large number of occurrences. Median EOO in these cases was as small as 12% of the actual value (inner 90% quantile range: 2-37%). Adding county records improved estimates in nearly every case and median estimates asymptotically approached “omniscient” EOO. Across all scenarios, EOO was overestimated in <2% of cases when county records were included.

**Figure 4.**
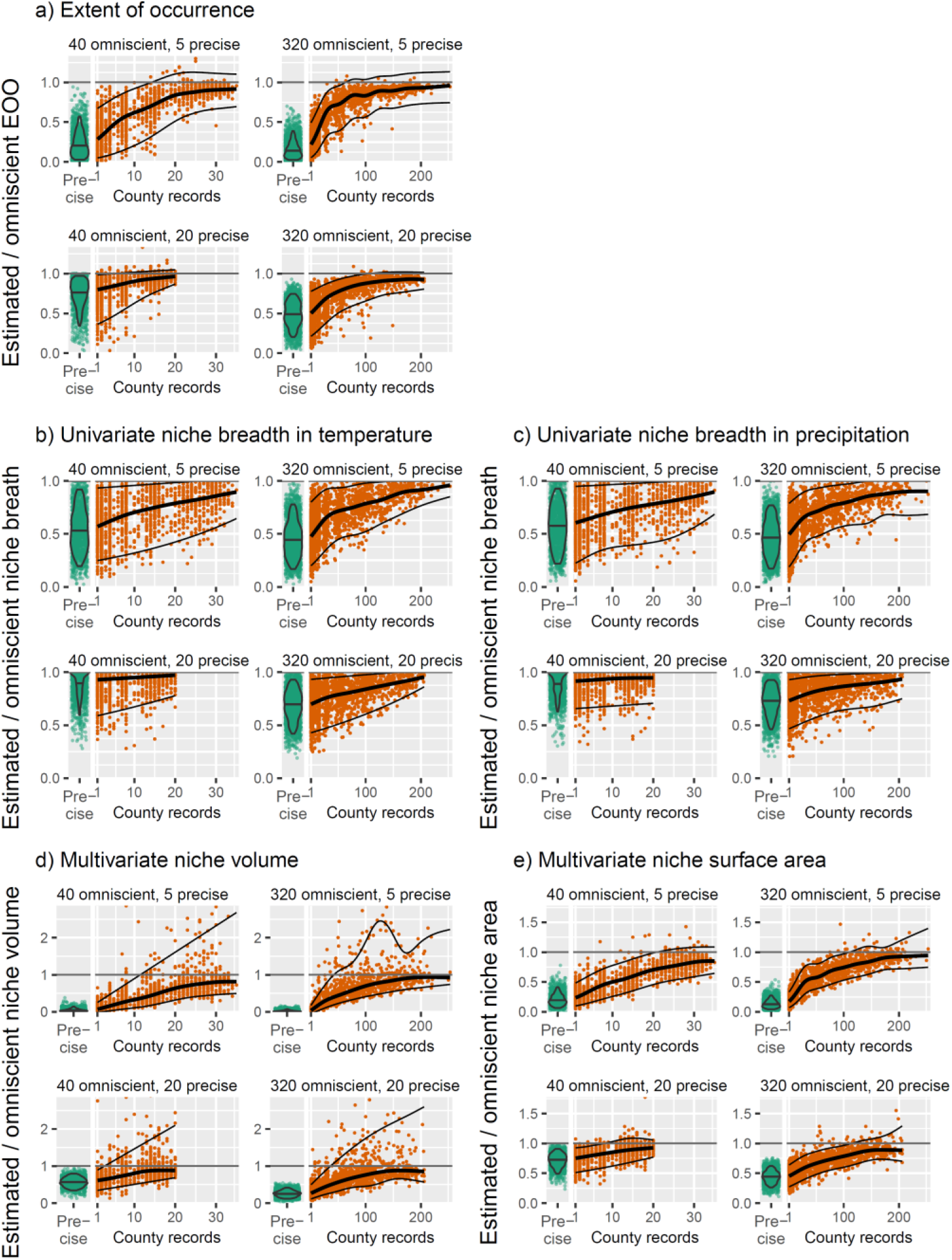
Adding imprecise records increases the accuracy of estimates of (a) extent of occurrence, (b) univariate niche breadth in mean annual temperature and (c) total annual precipitation, and (d) multivariate niche volume and (e) surface area. In each case, values represent the ratio of estimates using only precise or precise+imprecise records to estimates calculated using all “omniscient” occurrences of a species. Each green point represents a species analyzed using only precise records. Orange points represent these same species, but augmented with a particular number of imprecise records (indicated by the number of counties in which they occur). Violins encompass the inner 90% quantile of values. Thick trendlines represent the median trend, and thin trendlines encompass the inner 90% quantile of values. To aid visualization for univariate niche breadth and niche volume, the y-axis limits encompass only the lower 99.9% of values.

#### Niche breadth

Using just precise records underestimated univariate niche breadth and multivariate niche volume and surface area (Fig. 4b-e and Appendix S2 Figs. 2.8 to 2.11). For example, for a species with 40 total occurrences with 5 precisely-geolocated records, precise-only niche breadth in temperature was just 29% (6-71%) of omniscient niche breadth. However, precise+imprecise niche breadth in temperature increased up to ^~^90% (^~^70-110%) of omniscient breadth as county records were added. Estimates of niche breadth continued to improve for species with ≥20 precise occurrences, albeit at a slower rate as imprecise records were added. Similar trends were observed for precipitation, and in multivariate niche volume and surface area (Fig. 4c-d). Adding imprecise data led to occasional overestimation, especially for niche breadth in temperature and niche volume.

#### ENMs

Precise+imprecise models were more accurate than precise-only models when projected to both present and future climate scenarios (Figs. 5a and b, and Appendix 2 Figs. S2.1 and S2.2). Accuracy increased most when the number of precise records in precise+imprecise models was ≤15. For example, for present-day climate and a species with 40 total occurrences and only 5 precise occurrences (top left of panel Fig. 5a), the median correlation between the true probability of presence and precise-only ENM predictions was 0.15 (−0.02 to 0.58). Adding county records increased the correlation up to 0.63 (0.30-0.86), which was equivalent to the accuracy of omniscient ENMs (median: 0.60, 90% quantiles: 0.27-0.81). Similar results were obtained for species with a greater number of total occurrences and few precise records so long as the number of precise records was ≤^~^15. However, for species with ≥20 precise records, adding county-level records improved accuracy at a slower rate (Fig. 5a). Nonetheless, adding county records still increased accuracy (Fig. S2.1). Calibration accuracy decreased when ENMs were projected to the future, but precise+imprecise models were still more accurate than precise-only models when sufficient county records were included (Fig. 5b).

**Figure 5.**
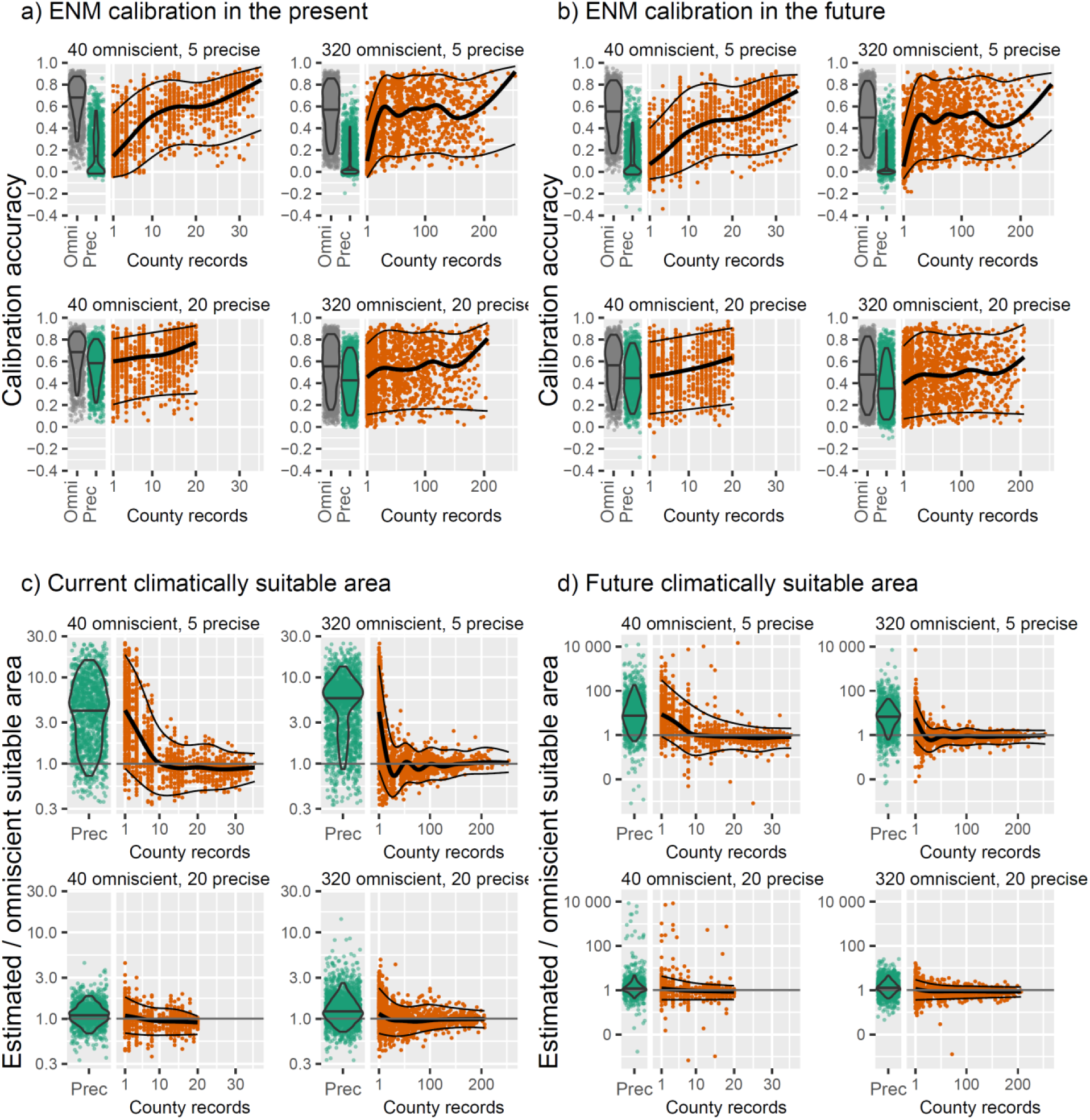
Adding imprecise records improves the accuracy of niche models for (a) the present and (b) future. Calibration accuracy is the correlation between predicted and actual environmental suitability. When only precise records are used, (c) current and (d) future climatically suitable area are overestimated, but including imprecise records reduces bias. Values are the ratio between estimates using precise-only or precise+imprecise records to estimates from models using all “omniscient” occurrences of a species. Each green point represents a species analyzed using only precise records. Orange points represent these same species, but augmented with a particular number of imprecise records (indicated by the number of counties in which they occur). Note the log scales. Violins encompass the inner 90% quantile of values. Thick trendlines represent the median trend, and thin trendlines encompass the inner 90% quantile of values. “Omni” refers to results from analyses using omniscient occurrences, and “Prec” to results using only precise occurrences.

Precise-only ENMs overestimated the area of climatically suitable habitat by up to >300% (median value) for the present (Fig. 5c) and up to >500% for the future (Fig. 5d and Appendix S.2 Figs. 2.6 and 2.7). Overestimation was greatest when the number of precise occurrences was <15. Adding just ^~^10-30 county-level records reduced bias to ^~^0 relative to omniscient models. Precise-only ENMs underestimated loss of suitable area by 90% or more when the number of precise records was ≤15 (Appendix S2 Fig. 2.3). Adding 10-30 county records eliminated this bias. Gains in suitable area were on average unbiased (Appendix S2 Fig. S2.4). Precise-only ENMs overestimated amount of area remaining climatically suitable through time (Appendix S2 Fig. 2.5). Adding imprecise records increased model complexity (Appendix S2 Fig. S13).

### Asclepias

The data obtained from GBIF comprised 53,623 herbarium records (Appendix S3 Table S3.1). Following standard data cleaning methods, removal of observational and duplicate records, and elimination of species with fewer than 5 geographically unique precise records, we were left with just 16% of the original records (8,480) and only 32% of the species (44 of 137). Imprecise records were the most abundant, comprising on average 70% of all usable records for a species (range: 44-97%). Thirty-six percent (16 of 44) of species had ≤20 precise records and twenty percent (9 of 44) had ≤10.

#### Extent of occurrence

Precise+imprecise EOO was 86% larger (median; range 0-2011%) than precise-only EOO (Fig. 6a). Including imprecise records at least doubled EOO for 34% of species (15 of 44), and at least tripled EOO for 27% of species (12 of 44). Adding imprecise records increased EOO more for species with fewer precise records.

**Figure 6.**
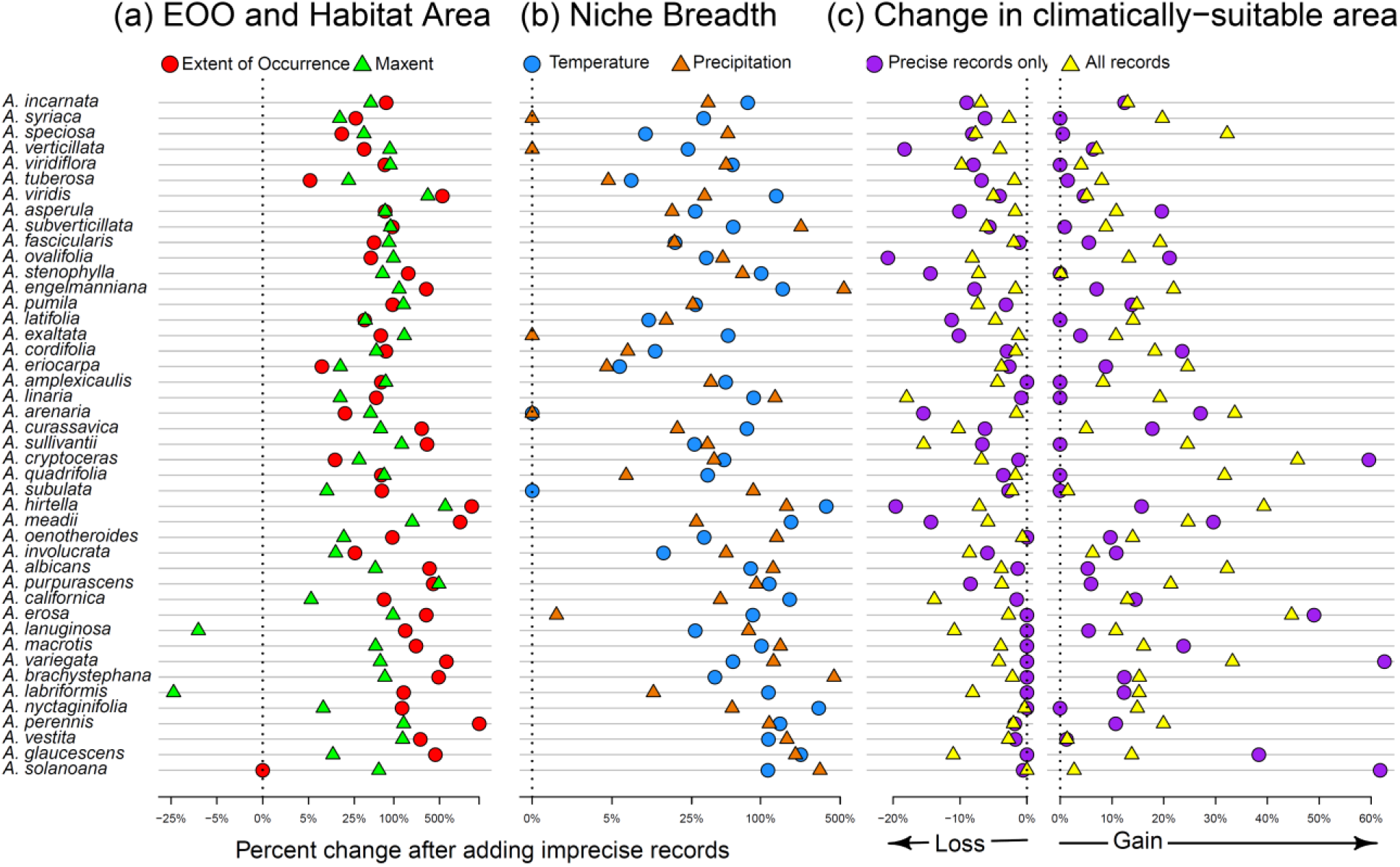
Analysis of North American *Asclepias* demonstrating the effects of including imprecise records on (a) EOO and present-day climatically suitable area predicted by ENMs; (b) univariate niche breadth in temperature and precipitation; and (c) gain and loss in climatically suitable area when niche models are predicted to the future. Ranges can both gain and lose area. Species are sorted from top to bottom from most to least number of precise records. Note the log scale along the x-axis.

#### Niche breadth

Including imprecise records increased median univariate niche breadth in temperature and precipitation by 25% (range: 0 to 353%) and 28% (0 to 292%) across species, respectively (Fig. 6b; Wilcoxon V ^~^0 and P<10^-6^ in both cases). Including imprecise records increased multivariate niche volume by a median value of 175% (8 to 13,909%; P<10^-12^, Wilcoxon V^~^0) and niche surface area by 79% (3 to 1515%; P<10^-12^, Wilcoxon V^~^0; Appendix S3 Figs. S3.3 and S3.4). Inclusion of imprecise records increased niche breadth, volume, and area most when species had few precise records.

#### ENMs

Relative to precise-only ENMs, precise+imprecise models predicted greater climatically suitable area for 63% of species (28 of 44) in the present and 71% of species (31 of 44) in the future (Fig. 6a and c). Gains and losses in suitable area were also larger when imprecise records were included (median gain 16% ranging from −80 to 3952%; median loss 30% ranging from −89 to 1652%; Appendix S3 Fig. S3.6).

## Discussion

Spatially imprecise occurrences are commonly discarded prior to biogeographic analyses (Fig. 1) to obviate propagation of spatial error through an analysis (Moudrý & Šímová 2012). However, discarding spatially imprecise records risks introducing error by undersampling geographic and environmental space. We found that including geospatially-imprecise records increased the accuracy of niche models projected to both present and future climate conditions (Fig. 5a and b). Including these records also led to more accurately estimated climatically suitable area (Fig. 5c and d) and improved estimates of niche breadth and extent of occurrence (EOO; Fig. 4). Sometimes estimates using just precise records were orders-of-magnitude different from those based on “omniscient” records, but adding imprecise records helped close this gap. Studies that have examined the effects of adding additional imprecise records to ENMs have typically not employed a conservative method for assigning environmental values or locations to imprecise records (e.g., Bloom et al. 2018; Collins et al. 2017; Cheng et al. 2021), or have focused on mean values (versus extremes, which define environmental limits; Pender et al. 2019). Our findings have direct implications for studies in ecology, evolution, and conservation that use occurrences to estimate species’ relationships to the environment.

### Why including imprecise records improves niche models and estimates of niche breadth

The effect of imprecise records on ENM accuracy and niche breadth depends on the abundance of precise records and how well they sample geographic and niche space (Moudrý & Šímová 2012; Tulowiecki et al. 2015; Soultan & Safi 2017). Including a sufficient number of imprecise records can fully compensate for a lack of precise records even precise records number as few as 5. Gains in accuracy were most notable for species with <^~^20 precise records, but even species >20 precise occurrences experienced improvements from adding imprecise records. Sample size is one of the largest influences on ENM accuracy (Santini et al. 2021), with minimum recommended sizes ranging from ^~^10 to several hundred (Wisz et al. 2008; van Proossdij et al. 2016; Santini et al. 2021; but see Rivers et al. 2011). Given that many species are known from just a handful of records (Zizka et al. 2018), including imprecise records could be especially helpful when sample sizes are small because they improve the sampling of geographic and environmental space (Fig. 4).

### Applicability of the NGP and NEP methods

The simplicity of the geometric basis behind the NGP and NEP procedures (Fig. 2) means that they are likely applicable in a wide variety of circumstances, including different geographic contexts like islands, different spatial resolution of predictors, different units like atlas occurrences, *et cetera*. These methods are particularly useful in undersampled regions. For example, consider *A. brachystephana*, for which only 8 precise locations were available (left panel in Fig. 2a). From the county- and municipios-level occurrences (middle panel of Fig. 2a), it is clear that its geographic distribution ranges much farther than precise records indicate. Precise records also undersample the climatic conditions it inhabits (Fig. 2b). In fact, NGP and NEP may be useful even in cases where all records are known “precisely.” After all, no georeferencing system—even GPS—is without error, even if small. Hence, even “precise” records have the chance to be erroneously matched to the wrong environmental cell if they fall close to cell borders. NGP and NEP are thus useful in a wide variety of situations.

Nonetheless, NGP and NEP may not be applicable or advisable in all situations. For example, categorical predictors cannot be used because there is no concept of “centroid” or “nearest” in qualitative space, although it may work for ordinal variables or numerical summaries of categories (e.g., percent forest cover). “Sample units” may also be too large to be used reliably with NEP. In particular, ENMs are sensitive not just to the values of variables but also their particular combination (Jiménez-Valverde et al. 2009; Mesgaran et al. 2014). Larger areas are likely to contain combinations of variables that are “novel” compared to environments accessible to a species. Thus, an environmental datum identified with NEP may be “nearest” to a given centroid, yet still possess combinations of variables very different from those preferred by a species. (For this reason, when analyzing *Asclepias*, we discarded imprecise records locatable only to areas larger than San Bernardino county, the largest “county” in North America; Appendix S3.). Finally, like precise records, imprecise records may be collected in a biased manner, requiring corrective modeling methods (Erickson & Smith 2021).

### Implications for studies in evolution, ecology, and conservation

Most species’ distributions remain poorly characterized despite enormous collection effort (Meyer et al. 2016). For *Asclepias*, even with >53,000 specimen records, we had enough data to analyze only a third species found in North America. Of those we could analyze, 70% of their records would have normally been discarded due to spatial uncertainty in their locations. Similar rates of spatial uncertainty are common in biodiversity databases (Moudrý & Devillers 2020).

Spatially imprecise records, if used carefully, have great potential to address the “Wallacean shortfall,” the lack of information on species’ distributions (Hortal et al. 2015), and the “Hutchinsonian” shortfall, the lack of information on species’ environmental tolerances (Cosentino & Maiorano 2021). Answers to many key questions in ecology and evolution are susceptible to these shortfalls, could gain from addition of erstwhile “unusable” imprecise records. For example, investigations focused on rates of climatic niche evolution (Saupe et al. 2018), niche breadth and range size (Quintero & Wiens 2013a and b), and measurements of niche overlap (Warren et al. 2008) will be inherently sensitive to the degree to which realized niches are adequately sampled. The many studies that rely on ENMs for reconstructing species’ past, present, and potential future distributions are especially sensitive to sample size (Santini et al. 2021) and uneven sampling intensity among inhabitable environments (Raes 2012; Perret & Sax 2022).

Discarding imprecise records can overestimate species’ vulnerability and thus bias conservation assessments. For example, under IUCN Red List criterion B1, species qualify as threatened if they have an EOO <100 km^2^ (in addition to other criteria; IUCN 2019; see also Young et al. 2016). While none of the 44 species of *Asclepias* in our analysis had an EOO <100 km^2^ when using just precise records, it is certainly possible that this threshold could be crossed by some of the other 93 species in our original data that we could not analyze because they had <5 precise records. Similarly, estimates of species’ adaptive capacities (Cang et al. 2016), rates of community thermophilization (Feeley et al. 2020), and exposure to anticipated climate change (Fig. 5d) could be misrepresented by undersampling due to removal of imprecise specimens.

Our intent is to provoke a reconsideration of the benefits and costs of discarding spatially imprecise records. These trade-offs must be assessed within the goals and philosophical approach of an analysis. Many conservation assessments adopt a precautionary strategy that errs on the side of assuming a species is more vulnerable than it may actually be (Moyle 2005; Huntley et al. 2016; IUCN 2019). In contrast, an evidentiary approach aims to classify species as vulnerable only if there is strong evidence to support such a designation (IUCN 2019). Discarding imprecise records decreased estimates of niche breadth and EOO (Fig. 4b-d), so aligns with a precautionary approach because species appear more vulnerable than they may be. However, using just precise records did overestimate climatically suitable area for the virtual species (Fig. 5c and d). This likely occurred because precise-only models were simpler (fewer coefficients; Appendix S2 Fig. S13), which tends to increase predicted suitable area (Brun et al. 2020). In contrast, EOO and niche breadth were occasionally overestimated when imprecise records were included (Fig. 4). Hence, whether or not a decision to retain versus keep imprecise records is precautionary or evidentiary depends on the metric used to assess vulnerability. When in doubt, we advocate capturing the full range of uncertainty by conducting precise-only with precise+imprecise analyses.

### Conclusions

We advocate a re-consideration of tradeoffs from discarding spatially imprecise occurrence records. Using only precise records reduces niche model accuracy, and can underestimate niche breadth and extent of occurrence. The decision over how to define imprecise records and whether or not to use them is an important contributor to the overall uncertainty in inherent in the outcome of analyses relying on specimen data. Discarding imprecise records ignores a critical aspect of uncertainty and risks undersampling species’ realized environmental and geographic distributions. Practitioners need to consider the trade-offs between using versus discarding imprecise records, especially given the preponderance of imprecise records available in specimen databases and the Wallacean and Hutchinsonian shortfalls that beset our knowledge of the distribution of life on Earth.

## Supporting information

Appendix S1: Literature survey of how imprecise records are handled in studies using SDMs

Appendix S2: Extended results for analyses using virtual species

Appendix S3: Extended methods and results for analyses using real species (Asclepias)

Appendix S4: ODMAP protocol report on modeling methods

## Acknowledgements

We thank Huijie Qiao, Neftalí Sillero, and an anonymous reviewer who helped us improve the manuscript, the Alan Graham Fund in Global Change and the Institute for Museum and Library Sciences (FAIN MG-30-15-0094-15) for supporting this work.

## Biosketch

The Global Change Conservation Lab at the Missouri Botanical Garden identifies solutions to pressing environmental problems by leveraging precise and the imprecise information on biodiversity. We are inspired by the cumulative person-millennia of field work and curatorial attention devoted to amassing collection records of Earth’s species and the exigency of using this data to its fullest potential.

## Data availability

Code for creating and analyzing the virtual species is available at https://github.com/adamlilith/enms_impreciseRecords. The climate data is available through the WorldClim portal (https://worldclim.org), and the data on *Asclepias* records was obtained from the Global Biodiversity Information Facility (doi: 10.15468/dl.uddn6o).

## Supplements

Appendix S1: Literature survey of how imprecise records are handled in studies using SDMs

Appendix S2: Extended results for analyses using virtual species

Appendix S3: Extended methods and results for analyses using real species (*Asclepias*)

Appendix S4: ODMAP protocol report on modeling methods

