## Appendix S1: Literature survey of how imprecise records are handled in studies using SDMs for "Including imprecisely georeferenced specimens improves accuracy of species distribution models and estimates of niche breadth"

[redacted for double-blind peer review]

### Appendix S1 Literature review of how spatially imprecise specimen records are handled in ecological niche modeling

We conducted a survey of the peer-reviewed species distribution modelling literature to determine common data cleaning practices. We were particularly interested in understanding common methods for dealing with imprecise records. A list of candidate articles was obtained using the same search criteria outlined by Araujo et al. (2019) who assessed common practices in distribution modeling, except that we restricted the results to articles written in English and published from 2010 to 2019. Articles were obtained using IS Web of Science (www.webofknowledge.com) on 2019-01-08.

The search resulted in 8470 total articles. Articles were first assessed for relevance by reading the titles and abstracts (Fig. S1). Articles were considered relevant if they mentioned using species distribution modeling, or synonyms or results thereof and used real (versus simulated) species. We then skimmed articles and flagged those that used museum or herbarium data in combination with some form of species distribution model, modeled real (versus virtual) species, and comprised the primary report of the modeling (i.e., versus reporting model results already appearing elsewhere in the published, peer-reviewed literature). Articles that used only data collected by the authors (regardless of whether specimens were ultimately deposited in a collection) and were excluded. We then scored these articles according to whether any data cleaning protocols were mentioned, whether or not records were discarded, and if so, the bases described for removing records. Bases included whether the records were:

- Indicative of individuals growing in “natural” conditions (e.g., versus being cultivated, purchased, etc.);
- Too spatially imprecise
- Geographic outliers (i.e., outside the species’ presumed range)
- Environmental outliers (i.e., outside the species’ expected niche)

We did not attempt to impose a pre-defined criterion for whether or not records were “unnatural”, too spatially imprecise, et cetera. Rather, we interpreted the authors’ statements tabula rosa and scored the article accordingly. Issues with spatial imprecision can be potentially ameliorated by high levels of spatial autocorrelation in the environmental variables (Moudrý & Šímová 2012) and/or increasing the grain size (resolution) of environmental layers (Graham et al. 2008). We did not score articles by whether or not they used larger cells to accommodate spatial imprecision in records because almost none provided a justification for the spatial resolution of the environmental data used in the analysis and none assessed the effect of spatial autocorrelation (vis-à-vis spatial imprecision) on the analysis (e.g., Naimi et al. 2014).

**Results**

Of 285 relevant articles, only 52% described any methods for cleaning data. Across publications that described data cleaning methods, 45% addressed issues related to coordinate uncertainty by removing records before the analysis (Fig. 1). Bases for removing records included being “unnatural” (e.g., cultivated or purchased), having imprecise or unknown coordinate uncertainty, lying outside the species’ presumed range, or falling outside the environmental tolerances of the species. No studies we reviewed used modeling methods that could account for uncertainty in spatial location or explicitly indicated that coarser resolution environmental data was used to accommodate spatially imprecise records.

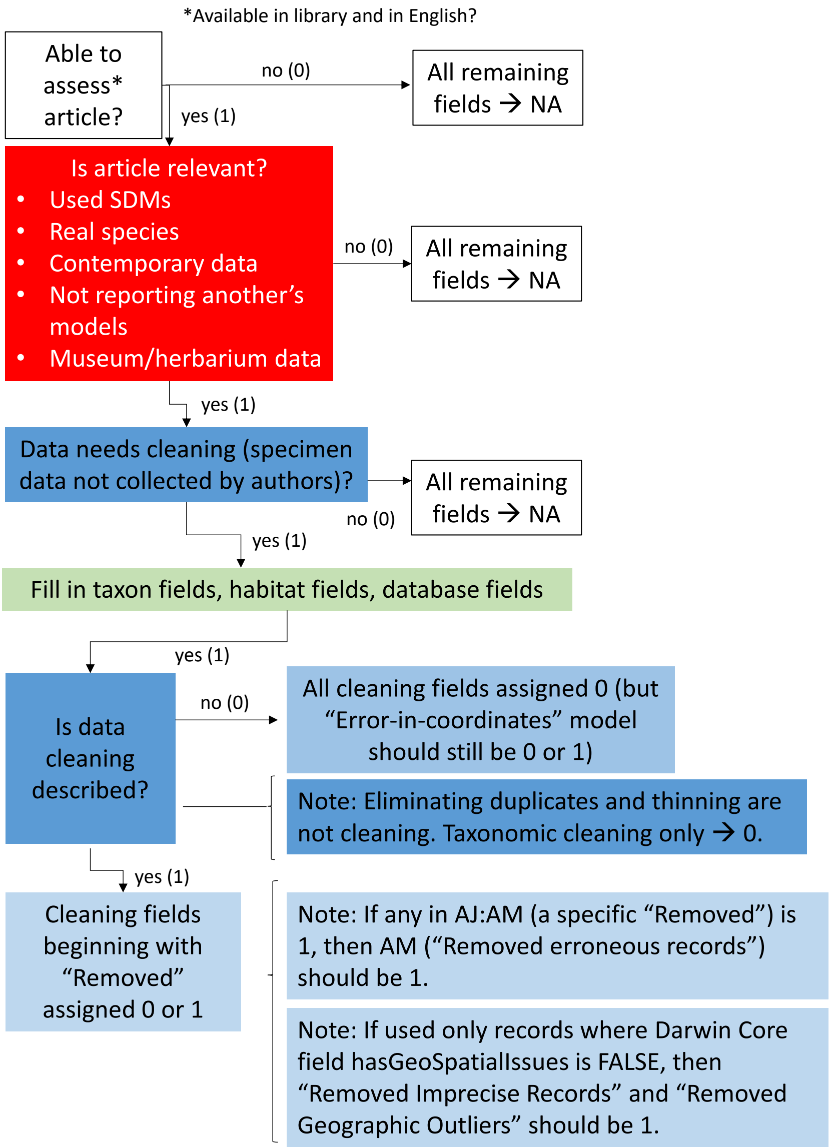


**Figure S1.1**. Decision trees for assessing data cleaning methods employed by articles using herbarium and/or natural history museum records and species distribution models.
