## Appendix S2: Extended results for analyses using virtual species for "Including imprecisely georeferenced specimens improves accuracy of species distribution models and estimates of niche breadth"

[redacted for double-blind peer review]

### Appendix S2 Extended results for virtual species

This supplement contains the full set of results for the analysis of virtual species.

**Figure S2.1** Calibration accuracy of ENMs against the true present-day probability of occurrence.

**Figure S2.2** Calibration accuracy of ENMs against the true future probability of occurrence.

**Figure S2.3** Loss in climatically suitable area estimated using ENMs.

**Figure S2.4** Gain in climatically suitable area estimated using ENMs.

**Figure S2.5** "Stable" climatically suitable area that remains suitable in time estimated using ENMs.

**Figure S2.6** Present-day climatically suitable area estimated using ENMs.

**Figure S2.7** Future climatically suitable area estimated using ENMs.

**Figure S2.8** Realized niche breadth in mean annual temperature (MAT).

**Figure S2.9** Realized niche breadth in total annual precipitation (TAP).

**Figure S2.10** Realized multivariate niche breadth volume.

**Figure S2.11** Realized multivariate niche surface area.

**Figure S2.12** Extent of occurrence (EOO).

**Fig. S2.13** Model complexity.

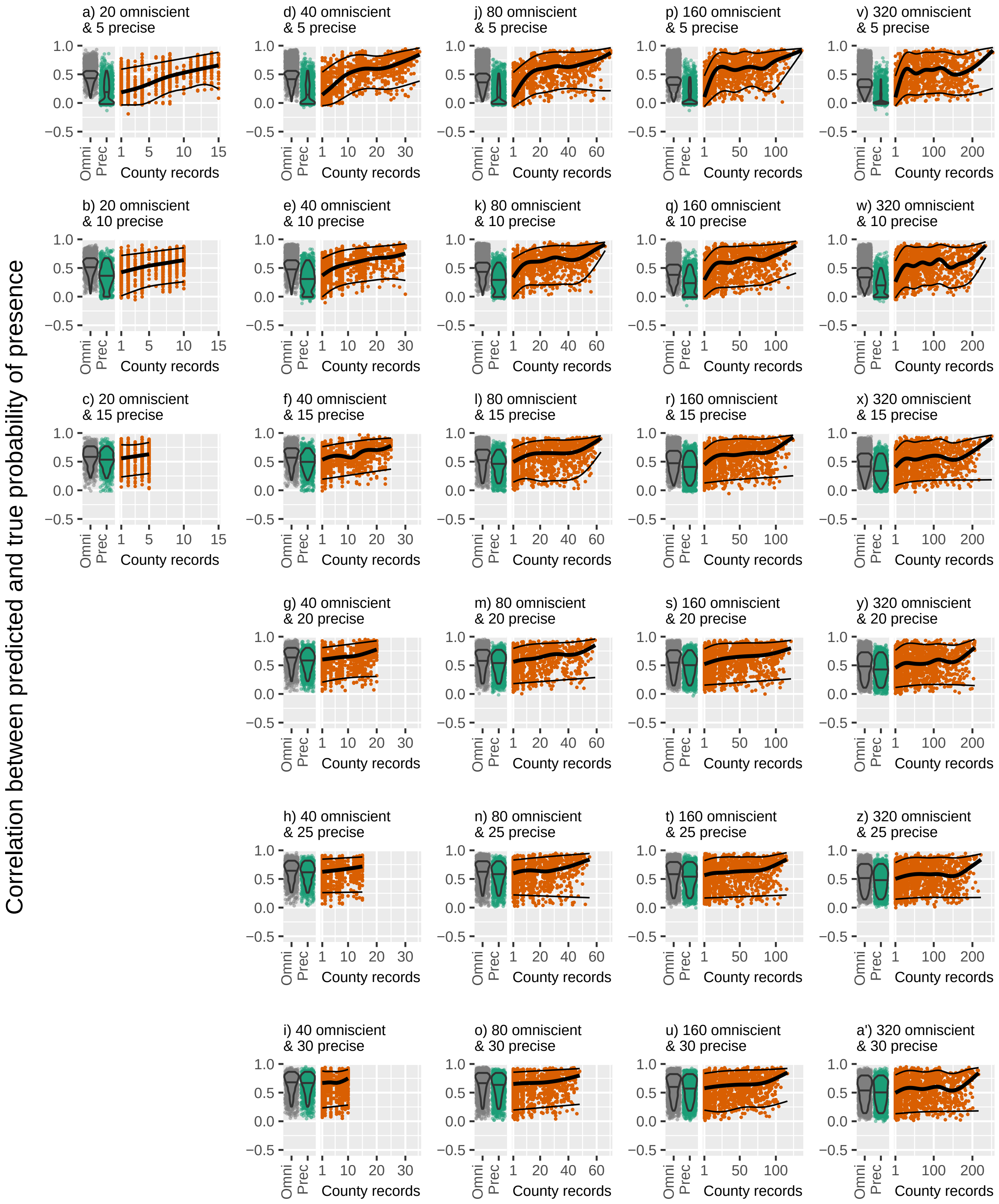


Figure S2.1 Calibration accuracy of ENMs against the true present−day probability of occurrence. Violin plots, and gray and green points represent the correlation between predictions from ENMs calibrated using omniscient and precise records (respectively) with the true probability of presence in the present. Green points represent the correlation between predictions from models calibrated using precise plus imprecise records and the true probability of presence. The thick black trend line traces the median trend and thin black lines bound the inner 90% of values. Higher values connote better calibration accuracy of ENMs.

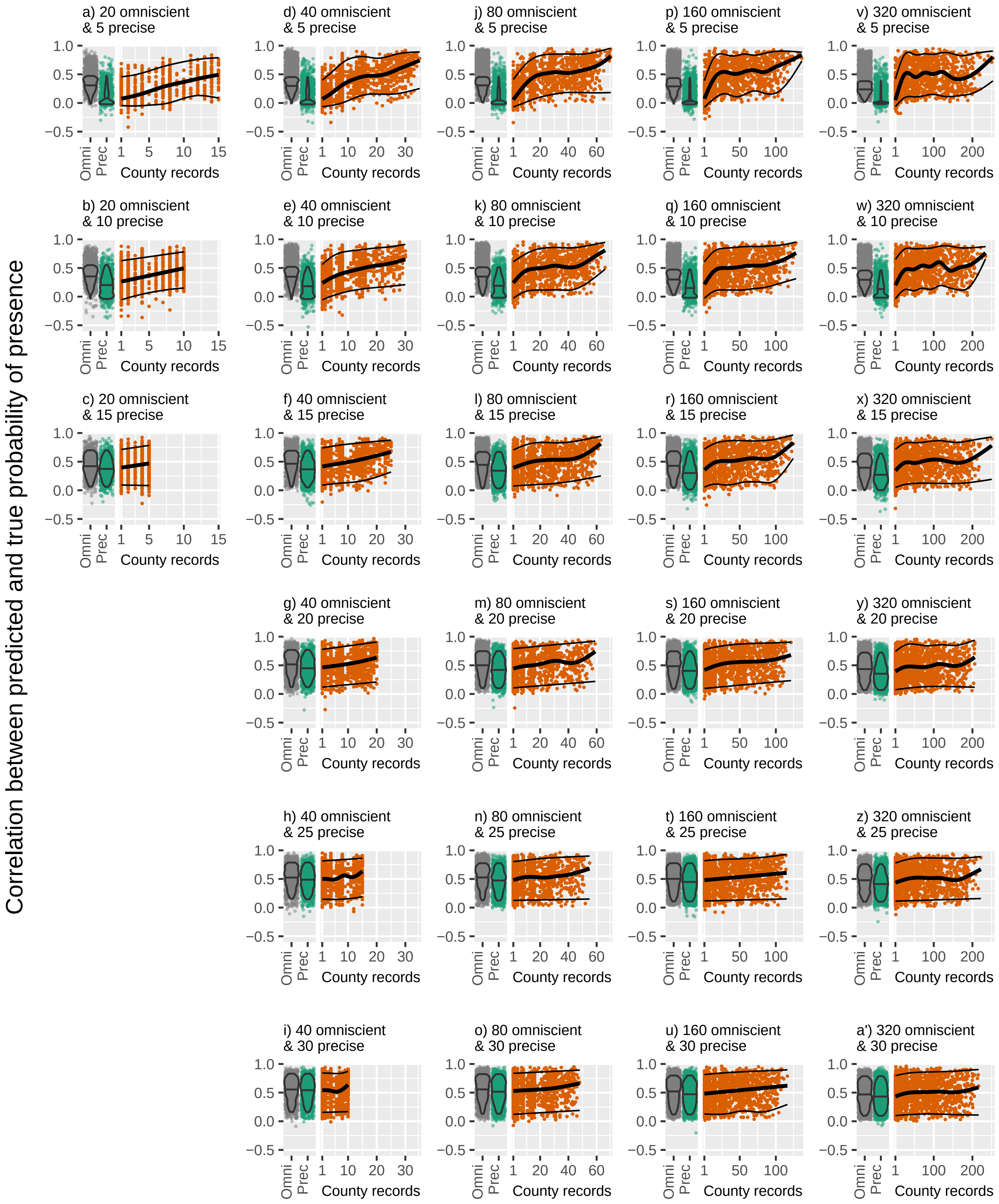


Figure S2.2 Calibration accuracy of ENMs against the true future−day probability of occurrence. Violin plots, and gray and green points represent the correlation between predictions from ENMs calibrated using omniscient and precise records (respectively) with the true probability of presence in the future. Green points represent the correlation between predictions from models calibrated using precise plus imprecise records and the true probability of presence. The thick black trend line traces the median trend and thin black lines bound the inner 90% of values. Higher values connote better calibration accuracy of ENMs.

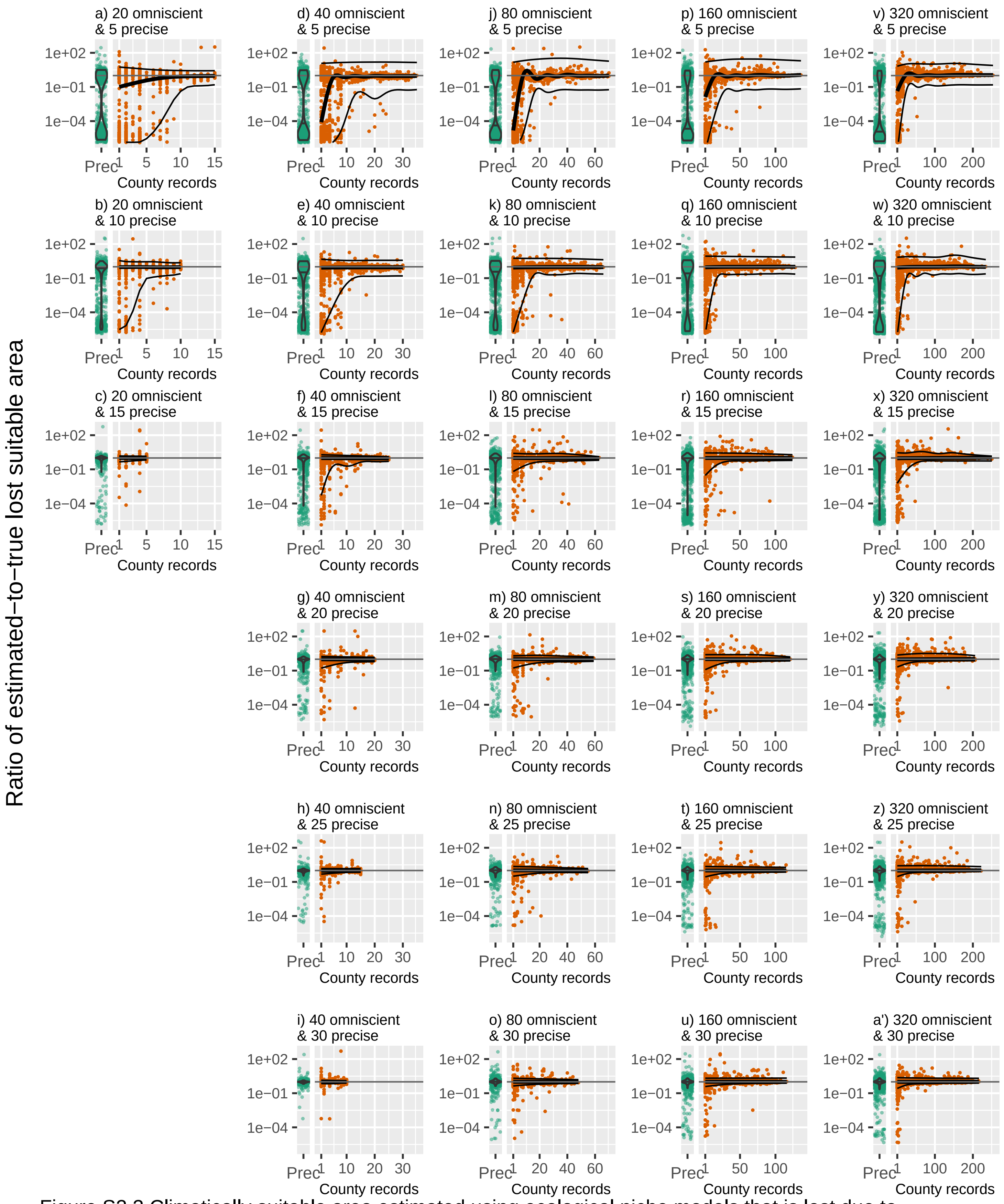


Figure S2.3 Climatically suitable area estimated using ecological niche models that is lost due to climate change. Violin plots and green points depict the ratio of lost suitable area estimated using only precise records to suitable area estimated using omniscient records. Orange points represent the ratio of lost suitable area estimated using precise plus imprecise records to lost suitable area estimated using omniscient records. The thick black trend line traces the median trend and thin black lines bound the inner 90% of values. Ratios were calculated as (estimated value + 1) / (omniscient value + 1). To aid visualization, only the inner 99.8% of values across all scenarios are shown.

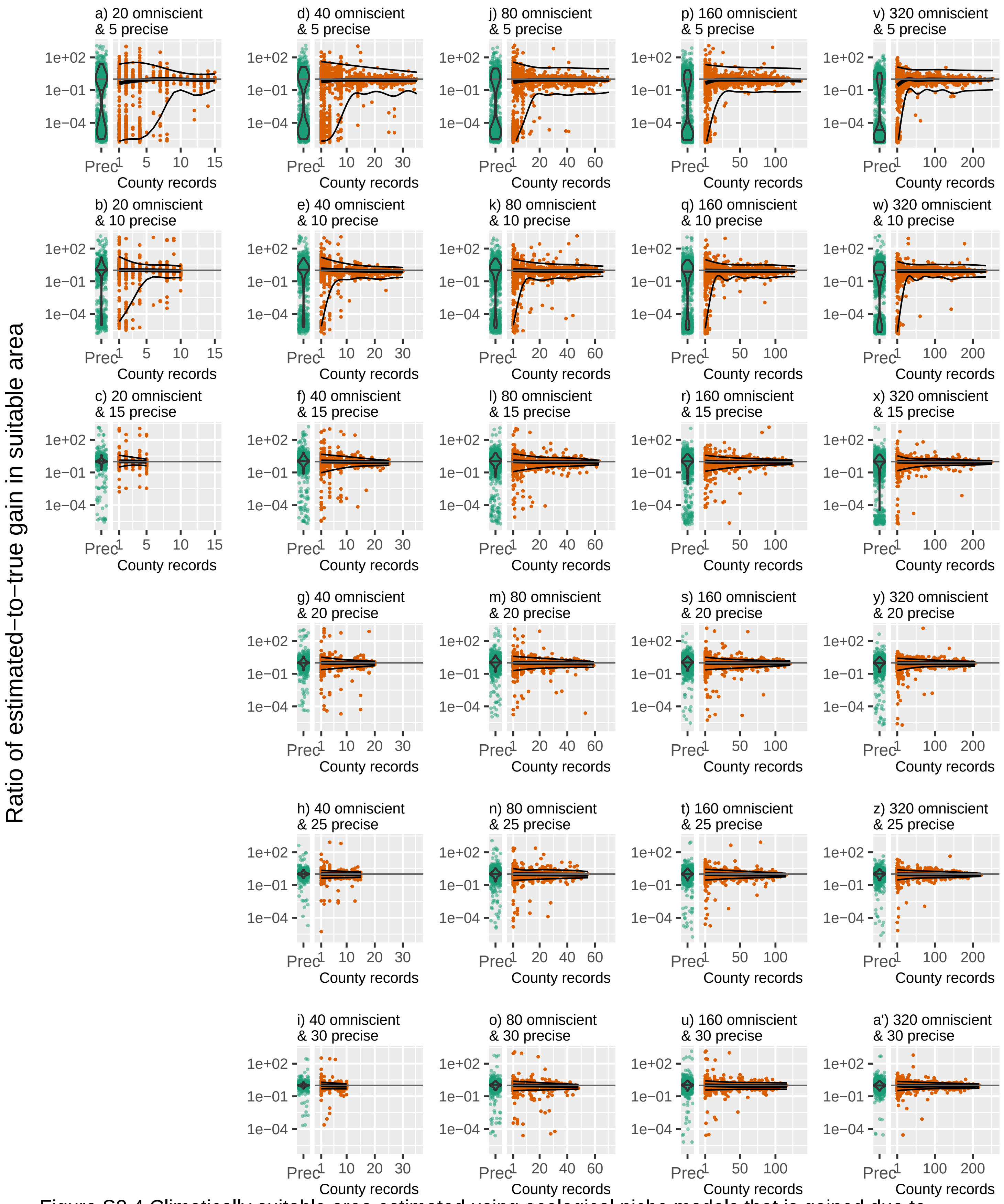


Figure S2.4 Climatically suitable area estimated using ecological niche models that is gained due to climate change. Violin plots and green points depict the ratio of gain in suitable area estimated using only precise records to gain in suitable area estimated using omniscient records. Orange points represent the ratio of gained suitable area estimated using precise plus imprecise records to gained suitable area estimated using omniscient records. The thick black trend line traces the median trend and thin black lines bound the inner 90% of values. Ratios were calculated as (estimated value + 1) / (omniscient value + 1). To aid visualization, only the inner 99.8% of values across all scenarios are shown.

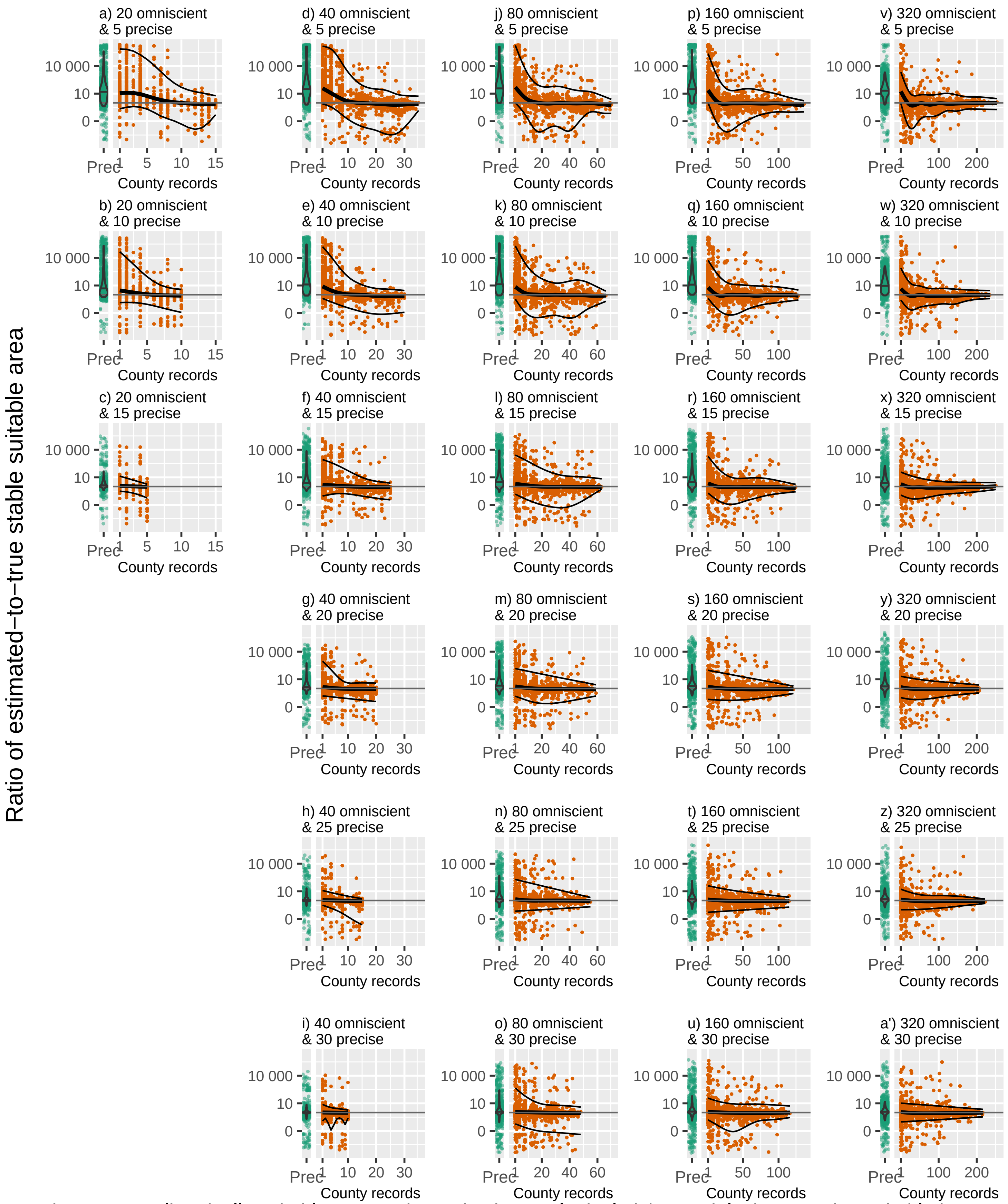


Figure S2.5 Climatically suitable area estimated using ecological niche models that remains suitable in time. Violin plots and green points depict the ratio of stable suitable area estimated using only precise records to suitable area estimated using omniscient records. Orange points represent the ratio of stable suitable area estimated using precise plus imprecise records to stable suitable area estimated using omniscient records. The thick black trend line traces the median trend and thin black lines bound the inner 90% of values. Ratios were calculated as (estimated value + 1) / (omniscient value + 1). To aid visualization, only the inner 99.8% of values across all scenarios are shown.

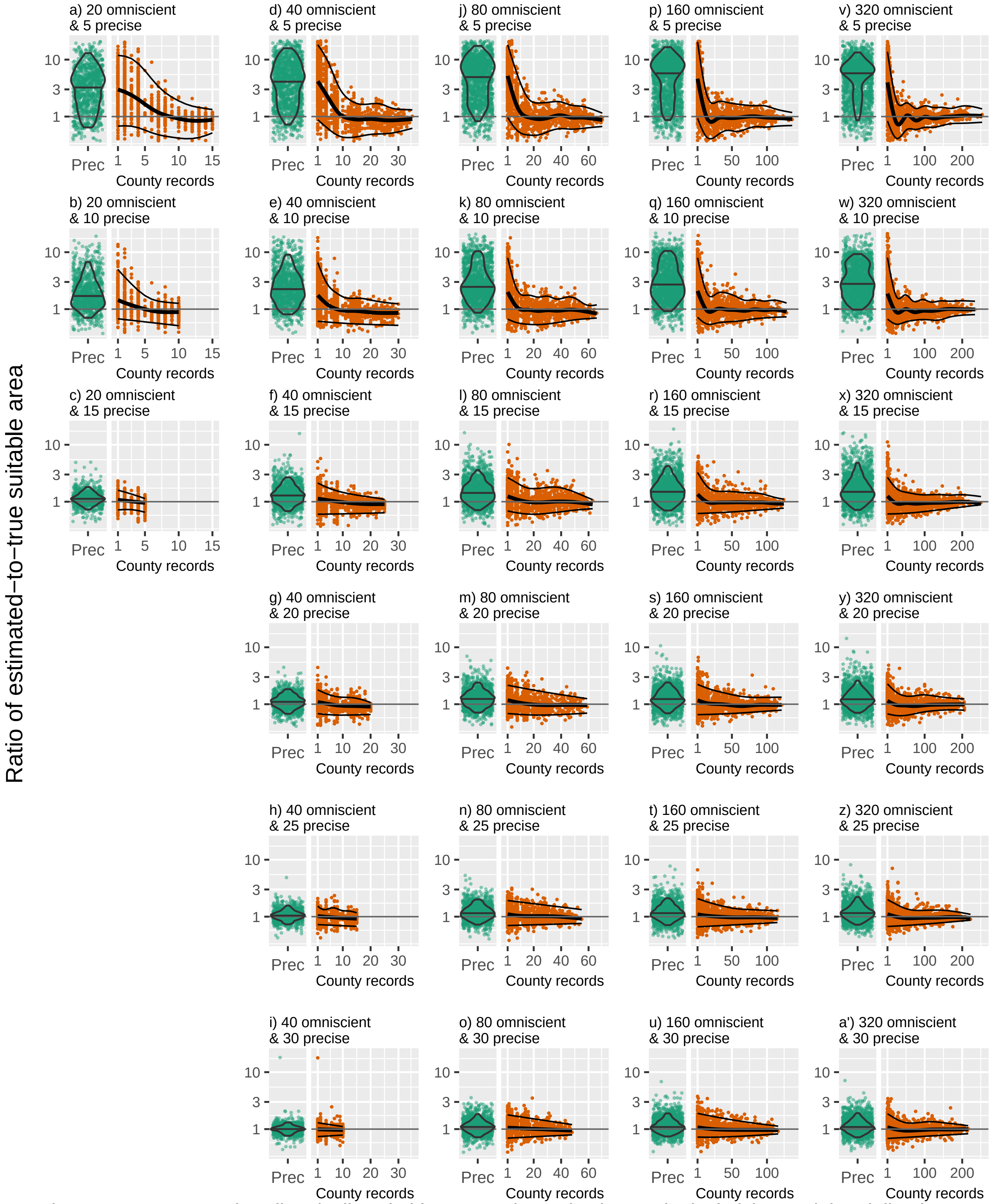


Figure S2.6 Present−day climatically suitable area estimated using ecological niche models. Violin plots and green points depict the ratio of present−day suitable area estimated using only precise records to suitable area estimated using omniscient records. Orange points represent the ratio of present−day suitable area estimated using precise plus imprecise records to suitable area estimated using omniscient records. The thick black trend line traces the median trend and thin black lines bound the inner 90% of values. To aid visualization, only the inner 99.8% of values across all scenarios are shown.

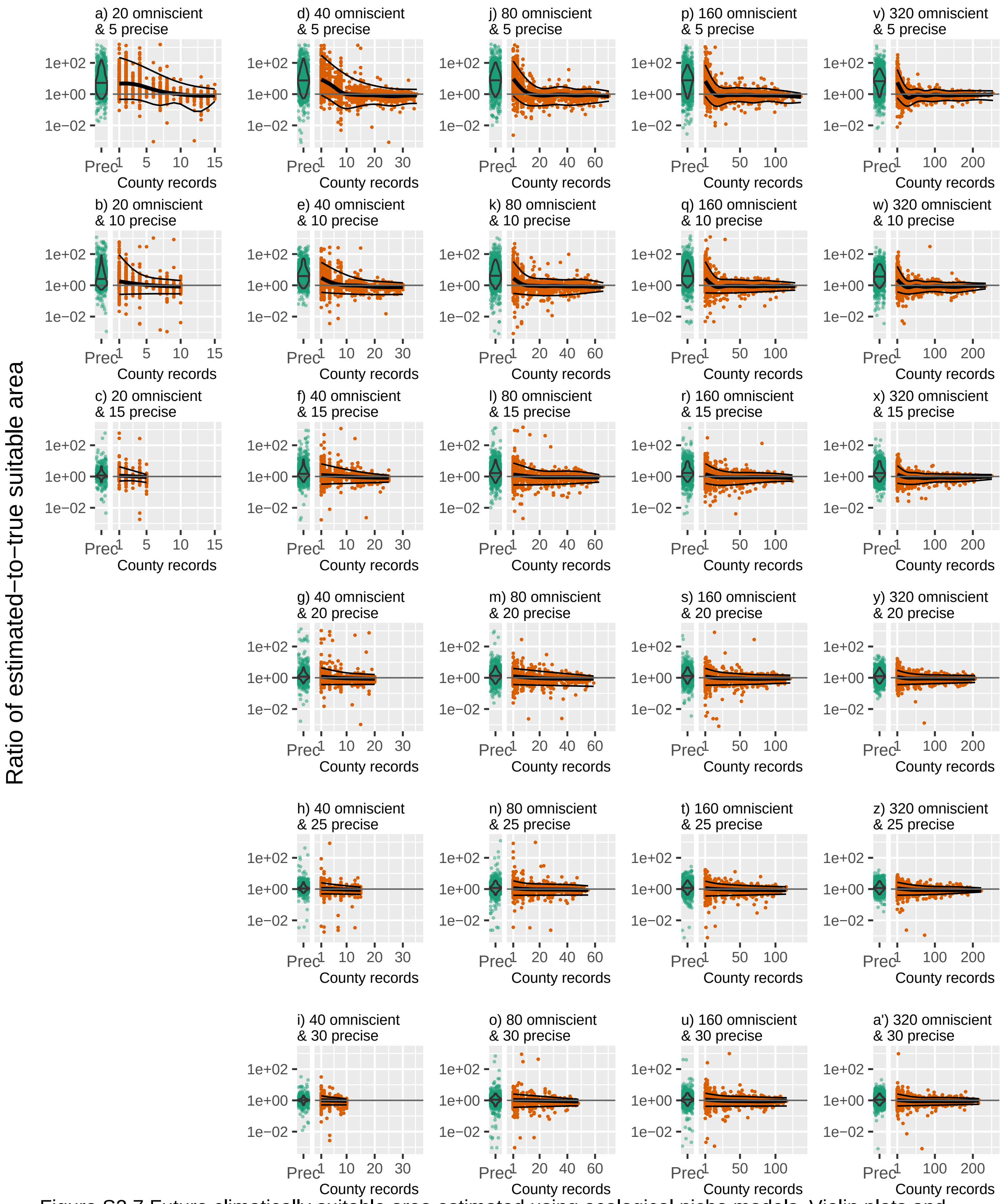


Figure S2.7 Future climatically suitable area estimated using ecological niche models. Violin plots and green points depict the ratio of future suitable area estimated using only precise records to suitable area estimated using omniscient records. Orange points represent the ratio of future suitable area estimated using precise plus imprecise records to suitable area estimated using omniscient records. The thick black trend line traces the median trend and thin black lines bound the inner 90% of values. Ratios were calculated as (estimated value + 1) / (omniscient value + 1). To aid visualization, only the inner 99.8% of values across all scenarios are shown.

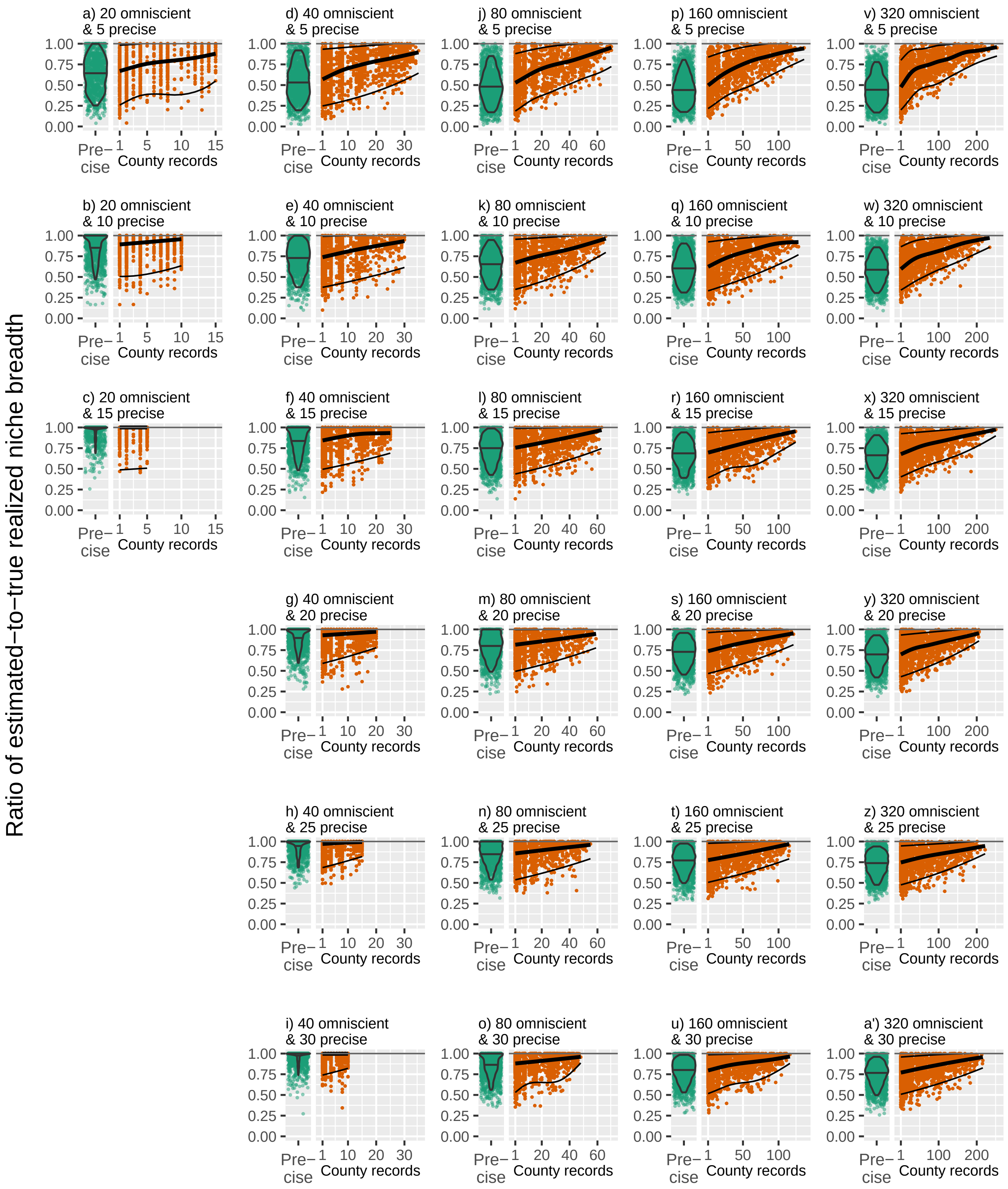


Figure S2.8 Estimates of realized niche breadth in mean annual temperature (MAT). In each panel the violin plot and green points show the ratio of realized niche breadth in MAT estimated using only precise records to the true realized niche breadth obtained from omniscient records. The violin plot encompasses the inner 90% of values and the horizontal line within represents the median value. The points and ratio of niche breadth estimated using precise plus imprecise records to true niche breadth (points). The thick black trend line traces the median trend and thin black lines bound the inner 90% of values. To assist in visualization, the top 0.1% of values across all scenarios are not shown.

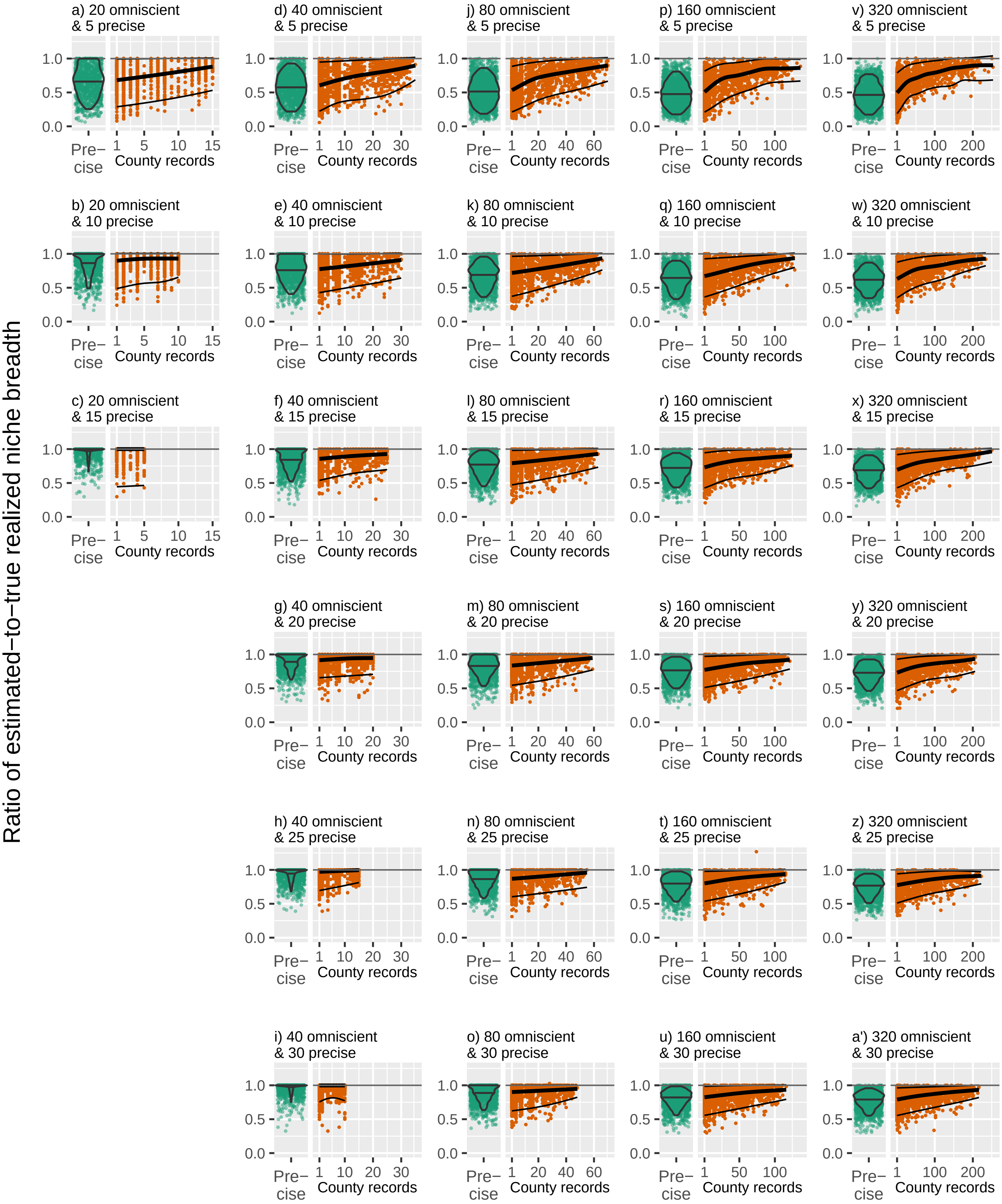


Figure S2.9 Estimates of realized niche breadth in total annual precipitation (TAP). In each panel the violin plot and green points show the ratio of realized niche breadth in TAP estimated using only precise records to the true realized niche breadth obtained from omniscient records. The violin plot encompasses the inner 90% of values and the horizontal line within represents the median value. The points and ratio of niche breadth estimated using precise plus imprecise records to true niche breadth (points). The thick black trend line traces the median trend and thin black lines bound the inner 90% of values.

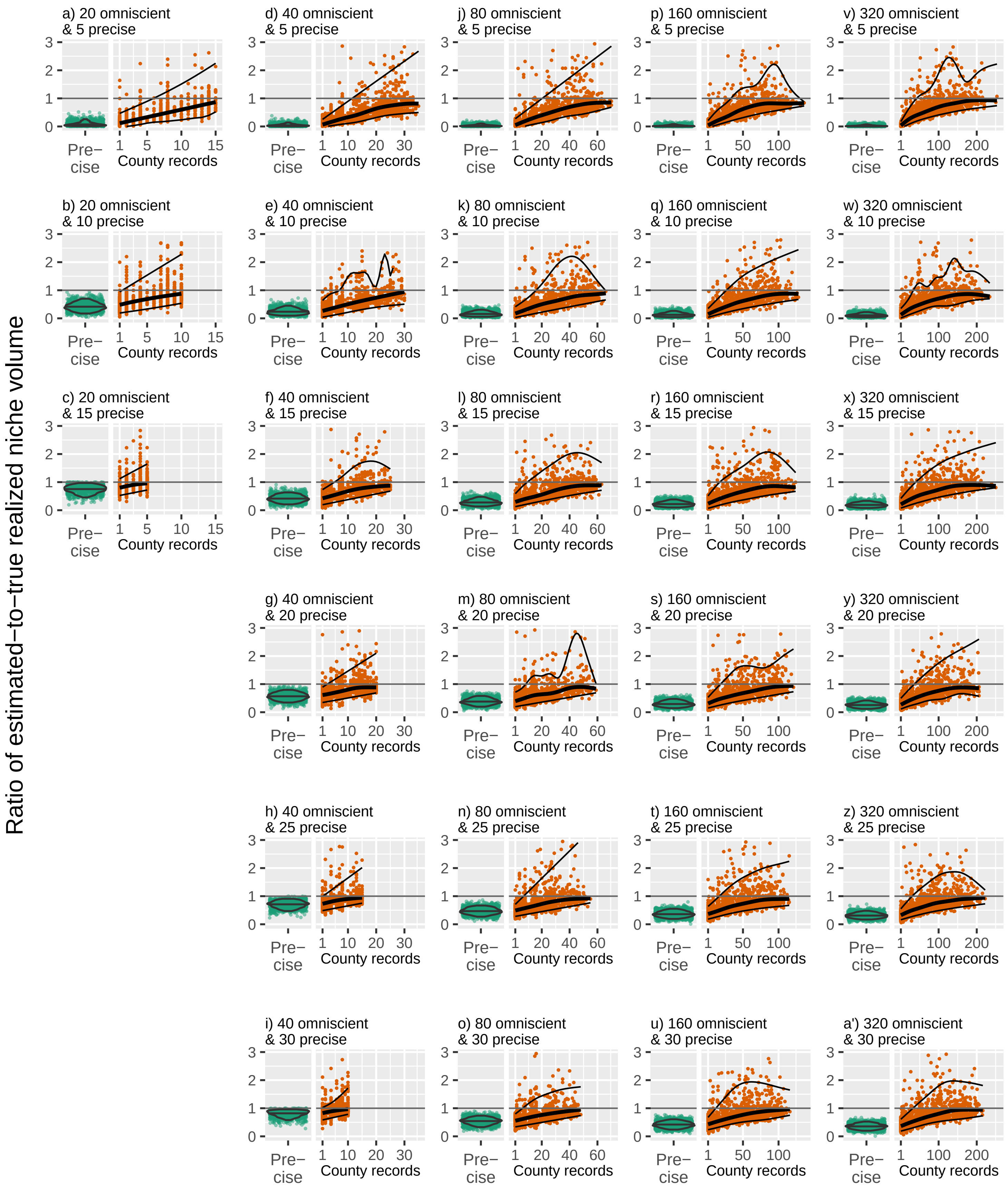


Figure S2.10 Estimates of realized multivariate niche breadth volume. The violin plot and green points show the ratio of realized niche volume estimated using only precise records to the true realized niche volume obtained from omniscient records. The violin plot encompasses the inner 90% of values and the horizontal line within represents the median value. The orange points represent the ratio of niche volume estimated using precise plus imprecise records to true niche breadth. The thick black trend line traces the median trend and thin black lines bound the inner 90% of values. To assist in visualization, the top 0.1% of values across all scenarios are not shown.

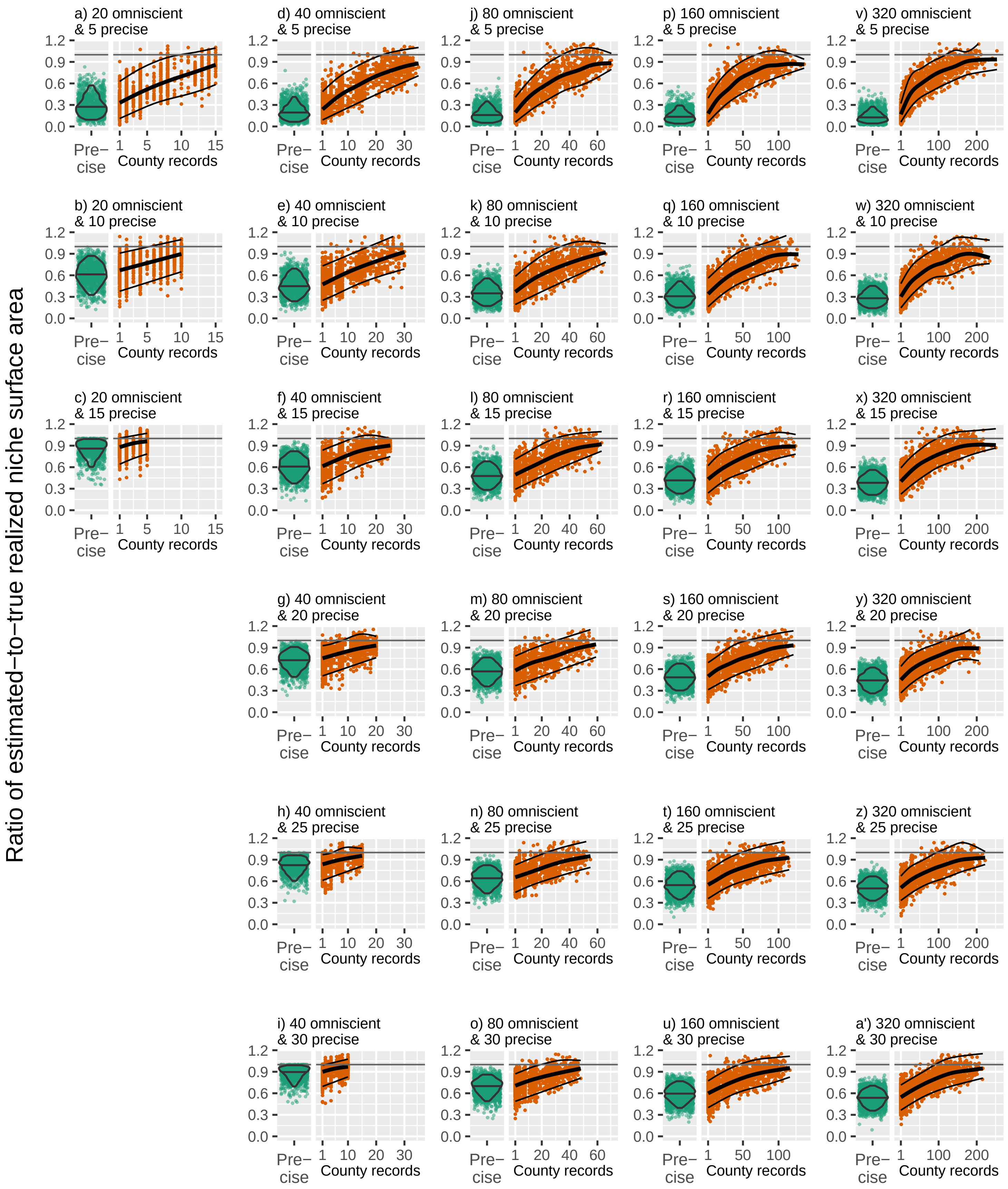


Figure S2.11 Estimates of realized multivariate niche surface area. The violin plot and green points show the ratio of realized niche area estimated using only precise records to the true realized niche area obtained from omniscient records. The violin plot encompasses the inner 90% of values and the horizontal line within represents the median value. The orange points represent the ratio of niche area estimated using precise plus imprecise records to true niche breadth. The thick black trend line traces the median trend and thin black lines bound the inner 90% of values. To assist in visualization, the top 0.1% of values across all scenarios are not shown.

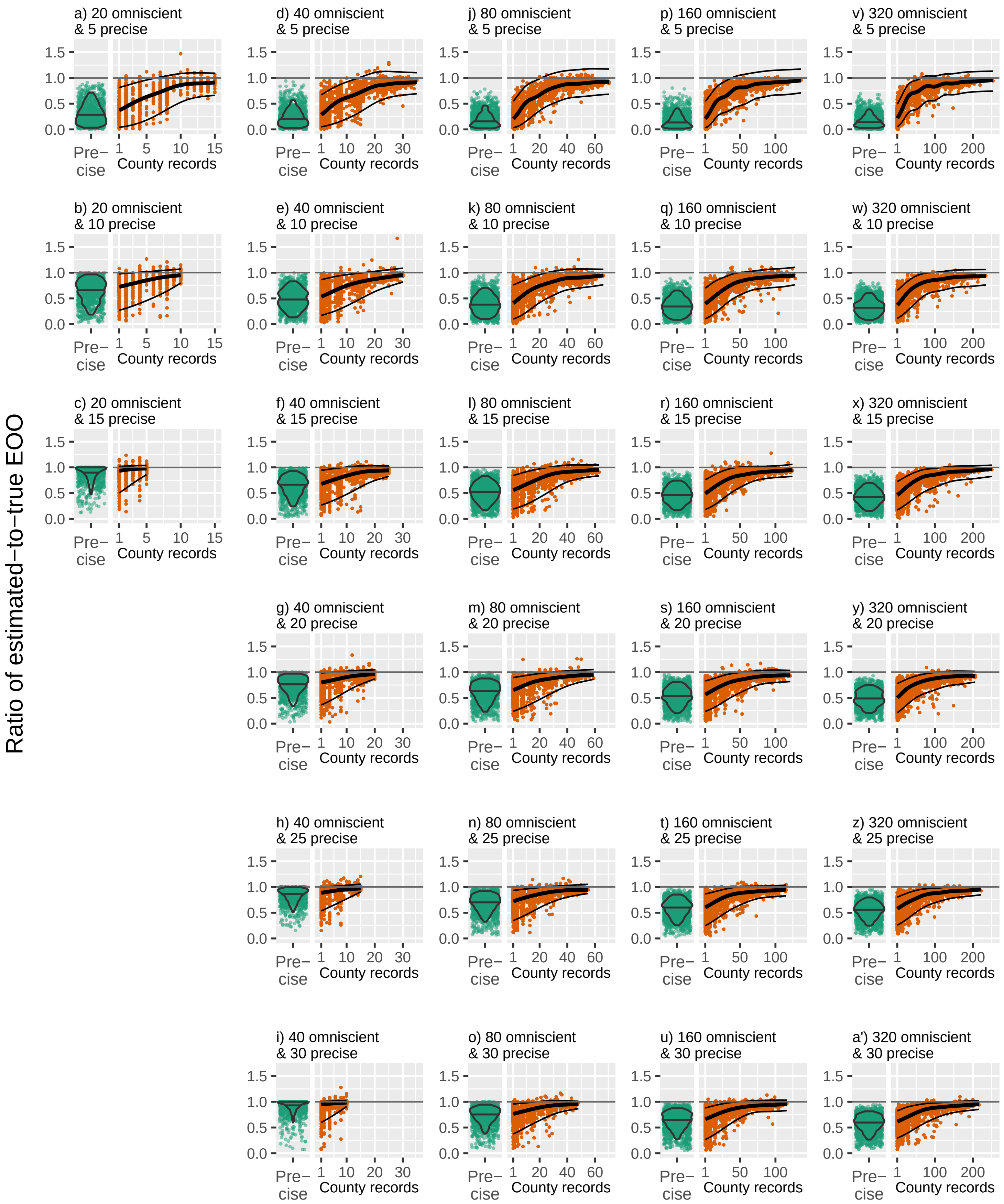


Figure S2.12 Estimates of extent of occurrence (EOO) compared to true EOO. The violin plot and green points depict the ratio of EOO estimated using only precise records to the true EOO obtained from omniscient records. The violin plot covers the inner 90% of estimates, and the median is represented by the horizontal bar within it. Orange points represent the ratio of EOO estimated using precise plus imprecise records to true EOO. The thick black trend line traces the median trend and thin black lines bound the inner 90% of values.

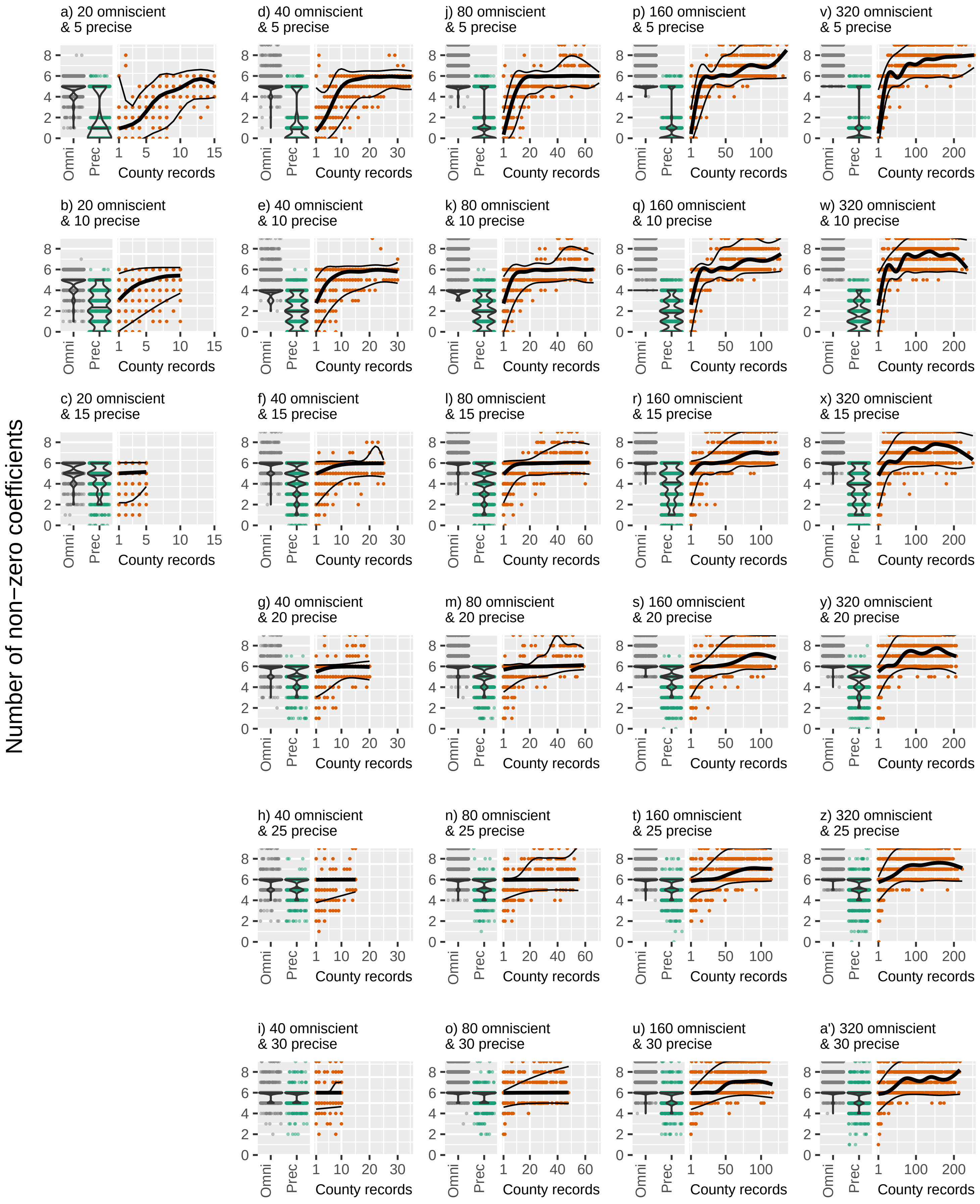


Figure S2.13 Model complexity measured as the number of non−zero coefficients, excluding the intercept. Violin plots, and gray and green points represent the number of coefficients of models using omniscient and precise records (respectively). Green points represent the number of coefficients of models using precise plus imprecise records. The thick black trend line traces the median trend and thin black lines bound the inner 90% of values. Higher values connote increased model complexity.
