## Appendix S3: Extended methods and results for analyses using real species (Asclepias) for "Including imprecisely georeferenced specimens improves accuracy of species distribution models and estimates of niche breadth"

**Including imprecisely georeferenced specimens improves accuracy of species distribution models and estimates of niche breadth: Don't let the perfect be the enemy of the good**

[redacted for double-blind peer review]

### Appendix S3 Methods and results for specimen cleaning and analysis of *Asclepias*

### Procedures for cleaning and classifying specimen records

We manually downloaded specimen data for *Asclepias* from the Global Biodiversity Information Facility (GBIF; REF) on November 18th, 2019 (doi: [redacted for double-blind peer review]). We restricted the search to records located in North America (Canada, the United States, and Mexico), but did not use any additional search filters within the GBIF search tool. After downloading, we removed records missing species names and records based on human observations (field "basisOfRecord" was any of "HUMAN_OBSERVATION", "OBSERVATION", or "MACHINE_OBSERVATION"). For records missing a value in the "eventDate" field, we used an automated routine to scrub the year of collection from the "verbatimEventDate" field. We then discarded records which could not be unambiguously assigned a collection year, or that were collected before 1970 to ensure an overlap with the temporal span of the climate coverages we used. We also removed records that were likely cultivated, purchased, or otherwise collected in conditions not indicative of natural environmental requirements by manually inspecting records identified using a search on the text in any of the "habitat", "locality" or "verbatiumLocalty" fields for strings "cultivat*", "grow*", "garden*", "experiment*", "captiv*", "greenhouse*", "arboretum", "hothouse*", "residential", "nursery*", "jardin*" (after forcing fields' text to lowercase). We manually cleaned names of states/provinces and counties to match those used in the Database of Global Administrative Areas (GADM ver 3.6; www.gadm.org; downloaded May 20, 2020). Then we removed records that could not be located to at least a state/province, that occurred outside the coterminous North America and minor outlying islands, or were collected in Hawaii.

Coordinate uncertainty is typically reported as the radius of a circle with a center at the given coordinate pair (Wieczorek et al. 2004), although not all records with coordinates report uncertainty (Moudrý & Devillers 2020). Uncertainty reflects imprecision in the method of geolocation (e.g., GPS versus maps-based interpolation), imprecision in the locality description (e.g., "northeast of Gettysburg" versus "7.3 miles northeast of Gettysburg on Old Harrisburg Road", and other aspects related to coordinate reference system and method of collection (Wieczorek et al. 2004). Coordinate precision reflects the number of significant digits in coordinate values (e.g., coordinates reported to 2 digits after the decimal can only be located with confidence to an area with a ~1.5-km radius at the equator; Meiri 2018). GBIF reports coordinates in decimal format, but coordinate precision may or may not be apparent from inspection of the coordinates in decimal format. For example, 38°37'37" is accurate to ±1 arcsec which is ±0.0002777° but could be reported as 38.626944, implying precision to ±0.000001°. Coordinate precision should be reflected in estimates of coordinate uncertainty (Wieczorek et al. 2004), but we found in some cases the uncertainty due to coordinate precision was larger than the stated uncertainty (no records had a value in the field "coordinatePrecision"). Additionally, some records had coordinates reported to a small number of decimal places but no associated coordinate uncertainty. To address this issue, we calculated coordinate uncertainty arising from coordinate imprecision based on the decimal and degree-minute-second representation of coordinates, used the larger of the two, then used either the stated coordinate uncertainty or uncertainty from imprecision, whichever larger, as the real coordinate uncertainty. Hereafter, we use "coordinate uncertainty" to refer to this value.

We categorized records into two groups, "precise" and "imprecise". Precise records were those with an associated coordinate uncertainty of ≤5000 m, which was about one-fourth the resolution of the environmental data we used (Graham et al. 2008). In some cases, the number of decimal places used to report coordinates implied a coordinate precision level that was less than the stated coordinate uncertainty due to georeferencing. In such cases, we used the larger of the stated uncertainty or uncertainty arising from truncated significant digits when categorizing records. In contrast, imprecise records comprised records that had a coordinate uncertainty ≥5000 m, or that could only be located to an administrative unit (county/parish or state/province). These latter records included those that were 1) missing coordinates but were able to be located to a geopolitical unit; 2) had coordinates but lacked coordinate uncertainty information but based on the number of significant digits in the coordinates could apparently be located to an area with a radius <10 km (assigned to a county) or ≥10 km (assigned to a state). Records that could not be assigned to either one of these categories were removed. We also discarded records containing conflicting information (e.g., noted as collected in a county that did not appear in the given state, or coordinates did not fall within the given county or state).We also removed all imprecise records that had an area of uncertainty larger than San Bernardino county, which is the largest "county"-level administrative unit in North America (52104.5 $km^2$) because we considered them too imprecise to convey salient information about the species' relationship to the environment.

Next, we created maps for each species and manually inspected records that seemed to fall outside the general vicinity of the majority of the records. Based on inspection of the data in the "habitat", "locality", "verbatimLocality" and other fields, we re-classified or removed these records. This allowed us to identify several records which were most likely geographic outliers or otherwise represented specimens from unnatural conditions (e.g., one georeferenced record was from a bridal bouquet).

Finally, we removed duplicate records of each species, which were defined based on the type of record. For precise records, we removed all but one record from each 10-arcmin cell in the rasterized climate surfaces that we used in the analysis (Fick & Hijmans 2016). We kept the record with the smallest coordinate uncertainty (or randomly selected, if tied). For records that could only be assigned to a geopolitical unit, we removed all but one randomly-selected record in that unit. For imprecise records with coordinates, we rounded coordinates to the third decimal place and rounded coordinate uncertainty to the nearest 1000 m. We then designated records with the same rounded values as duplicates, and removed all but one randomly-selected duplicate. When finished, we created a second set of maps for each species and double-checked any records which appeared to be geographic outliers. Finally, we removed species for which we had fewer than 5 non-duplicated precise records, resulting in 44 species for the analysis (Table S1).

**Table S3.1**. Number of records of each type for each species after data cleaning, removing duplicated records, and discarding species with fewer than 5 precise records.

| | | Imprecise | | | |
|---|---|---|---|---|---|
| Species | Precise | Point-radius | Geopolitical | Subtotal | Imprecise / Total |
| *Asclepias albicans* | 17 | 6 | 22 | 28 | 0.62 |
| *Asclepias amplexicaulis* | 27 | 28 | 154 | 182 | 0.87 |
| *Asclepias arenaria* | 25 | 19 | 28 | 47 | 0.65 |
| *Asclepias asperula* | 77 | 44 | 143 | 187 | 0.71 |
| *Asclepias brachystephana* | 8 | 13 | 72 | 85 | 0.91 |
| *Asclepias californica* | 14 | 0 | 18 | 18 | 0.56 |
| *Asclepias cordifolia* | 32 | 2 | 32 | 34 | 0.52 |
| *Asclepias cryptoceras* | 24 | 2 | 44 | 46 | 0.66 |
| *Asclepias curassavica* | 25 | 9 | 477 | 486 | 0.95 |
| *Asclepias engelmanniana* | 51 | 31 | 101 | 132 | 0.72 |
| *Asclepias eriocarpa* | 29 | 1 | 22 | 23 | 0.44 |
| *Asclepias erosa* | 12 | 4 | 28 | 32 | 0.73 |
| *Asclepias exaltata* | 38 | 7 | 102 | 109 | 0.74 |
| *Asclepias fascicularis* | 69 | 7 | 109 | 116 | 0.63 |
| *Asclepias glaucescens* | 5 | 0 | 144 | 144 | 0.97 |
| *Asclepias hirtella* | 21 | 11 | 78 | 89 | 0.81 |
| *Asclepias incarnata* | 239 | 101 | 519 | 620 | 0.72 |
| *Asclepias involucrata* | 19 | 6 | 28 | 34 | 0.64 |
| *Asclepias labriformis* | 7 | 0 | 6 | 6 | 0.46 |
| *Asclepias lanuginosa* | 12 | 11 | 25 | 36 | 0.75 |
| *Asclepias latifolia* | 40 | 43 | 42 | 85 | 0.68 |

| | | | | | |
|---|---|---|---|---|---|
| *Asclepias linaria* | 27 | 19 | 343 | 362 | 0.93 |
| *Asclepias macrotis* | 10 | 1 | 31 | 32 | 0.76 |
| *Asclepias meadii* | 21 | 2 | 24 | 26 | 0.55 |
| *Asclepias nyctaginifolia* | 6 | 1 | 15 | 16 | 0.73 |
| *Asclepias oenotheroides* | 20 | 8 | 168 | 176 | 0.90 |
| *Asclepias ovalifolia* | 68 | 48 | 36 | 84 | 0.55 |
| *Asclepias perennis* | 6 | 2 | 135 | 137 | 0.96 |
| *Asclepias pumila* | 41 | 46 | 35 | 81 | 0.66 |
| *Asclepias purpurascens* | 15 | 13 | 122 | 135 | 0.90 |
| *Asclepias quadrifolia* | 24 | 7 | 146 | 153 | 0.86 |
| *Asclepias solanoana* | 5 | 1 | 7 | 8 | 0.62 |
| *Asclepias speciosa* | 176 | 86 | 168 | 254 | 0.59 |
| *Asclepias stenophylla* | 52 | 14 | 57 | 71 | 0.58 |
| *Asclepias subulata* | 23 | 8 | 27 | 35 | 0.60 |
| *Asclepias subverticillata* | 70 | 20 | 106 | 126 | 0.64 |
| *Asclepias sullivantii* | 25 | 12 | 66 | 78 | 0.76 |
| *Asclepias syriaca* | 192 | 56 | 380 | 436 | 0.69 |
| *Asclepias tuberosa* | 151 | 41 | 512 | 553 | 0.79 |
| *Asclepias variegata* | 9 | 4 | 157 | 161 | 0.95 |
| *Asclepias verticillata* | 167 | 65 | 335 | 400 | 0.71 |
| *Asclepias vestita* | 6 | 1 | 11 | 12 | 0.67 |
| *Asclepias viridiflora* | 154 | 71 | 209 | 280 | 0.65 |
| *Asclepias viridis* | 80 | 18 | 168 | 186 | 0.70 |
| Median | 25 | 10 | 75 | 87 | 0.70 |
| Mean | 48.6 | 20.2 | 123.9 | 144.1 | 0.72 |
| Minimum | 5 | 0 | 6 | 6 | 0.44 |
| Maximum | 239 | 101 | 519 | 620 | 0.97 |

### Methods for ecological niche modeling of *Asclepias*

**Models using only precise records:**

Assigned climate values to records from cells in which each record fell

**Models using precise and imprecise records:**

Assigned climate values to precise records from cells in which each record fell

Assigned climate values to imprecise records in four ways:

- Value closest to mean across precise records' climate
- Value farthest from mean across precise records' climate
- Mean value within locality of likely collection
- Value at centroid of locality of likely collection

**All models:**

Used two definitions of background region:

- 300-km buffer around each occurrence (precise-only or precise & imprecise)
- 300-km buffer around minimum convex polygon around all records (precise-only or precise & imprecise)

Projected to 2070s assuming:

- Low-warning scenario (RCP 4.5)
- High-warming scenario (RCP 8.5)

**Figure S3.1** Outline of the niche modeling procedures for estimating exposure to climate change.

We used ecological niche models to estimate exposure to anticipated climate change. Please also see the ODMAP protocol for further details (Appendix S4; Zurell et al. 2020). Models were calibrated using the first 3 axes of the PCA. Background data were obtained from either an area defined by a 300-km buffer around the MCP of all occurrences, or from an area defined by a 300-km buffer around each occurrence. The former method assumes a species can disperse to all intermediate locations between known occurrences, but potentially overestimates the total area to which a species could disperse when the range is disjunct, while the latter obviates issues with disjunct distributions, but is more sensitive to the density of samples across the range. Separate buffers were used for models using only precise records and for all records combined. We assigned environmental values to imprecise records using the same methods described above for calculating niche breadth.

We modeled species' niches using Maxent ver. 3.3.3k (Phillips et al. 2006; Phillips & Dudík 2008) using precise and precise plus imprecise records. We calibrated Maxent models with linear, quadratic, and product terms and set the optimum master regularization parameter by exploring all possible combinations of terms plus values of the regularization parameter

(0.5, 1, 1.5, …, 5, 7.5, 10), then selecting the model with the lowest $AIC_c$ (Warren & Siefert 2011). All told, for each species we constructed 10 models (for models using only precise records: 2 background definitions; for models using precise and imprecise records: 2 background definitions x 4 methods for assigning climate variables to imprecise records). Each model was projected to a present-day scenario and two future emissions scenarios, RCPs 4.5 and 8.5. We thresholded present-day and future model output using a training specificity (true positive rate) rate of 90%. To estimate the current climatically suitable area, we calculated the percentage of the background region that fell above this threshold. To estimate exposure to climate change, we calculated the percentage of the present-day area predicted to be occupied that experienced gains and losses under future emissions scenarios (a species can experience an increase in suitable area in one place and a decrease in another). Comparisons between models with and without imprecise records were conducted within each combination of background (convex hull, circular buffers) and emissions scenario.

### Results

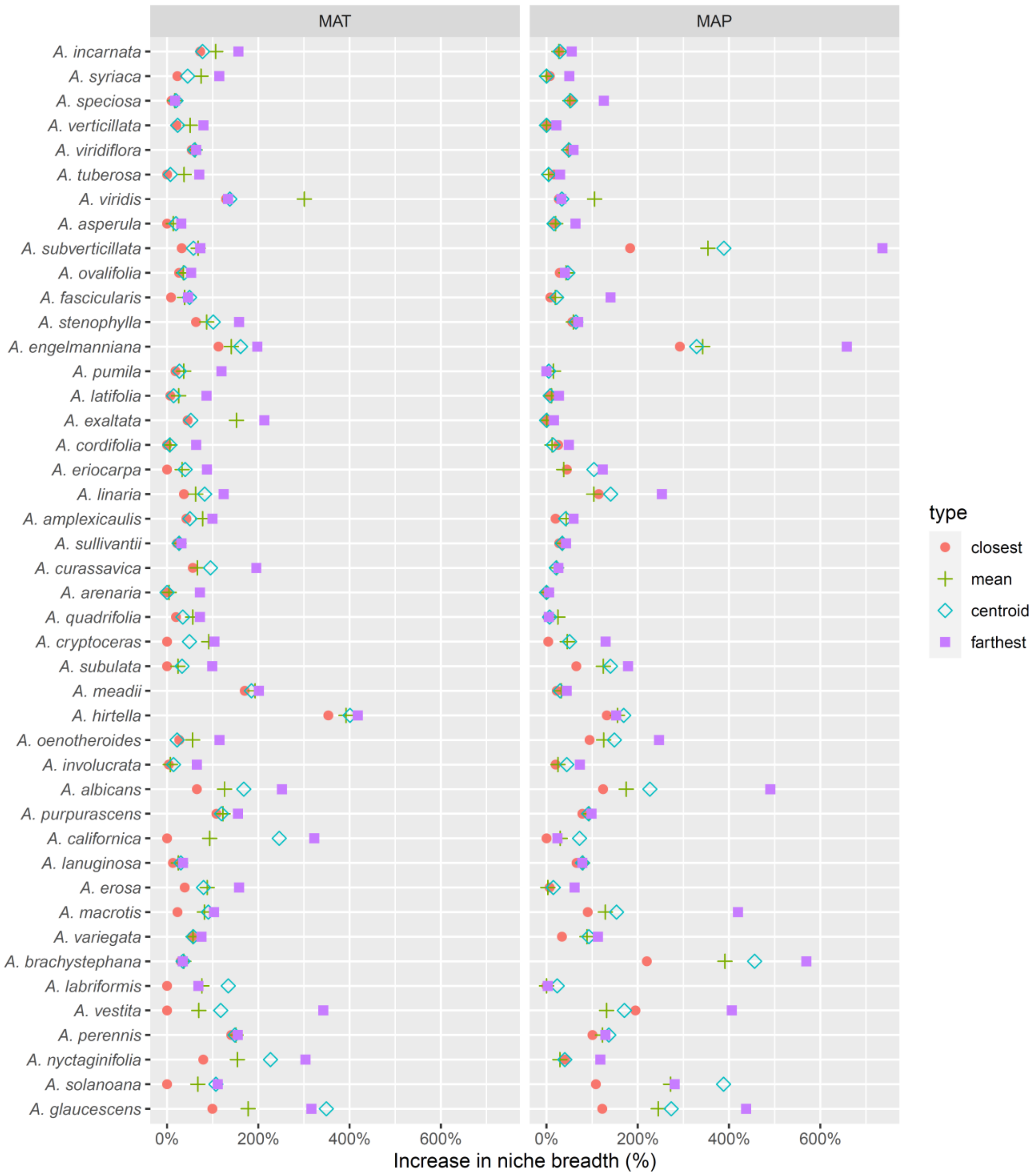


**Figure S3.2** Increase in **univariate niche breadth** in mean annual temperature (left) and precipitation (right) when imprecise records are included. Species are sorted from most to least number of precise records (same order as in Fig. 6).

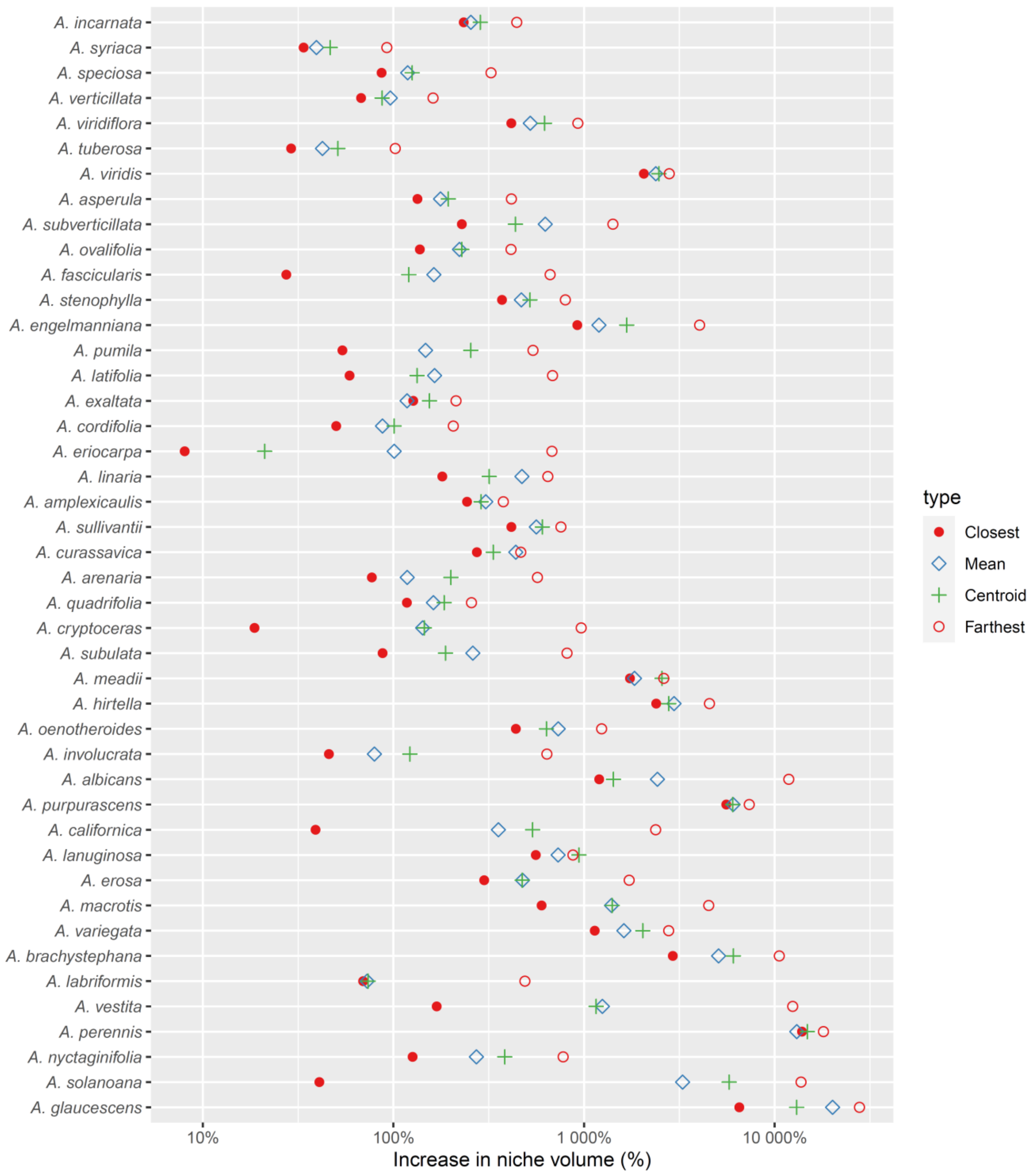


**Figure S3.3** Increase in **multivariate niche volume** when imprecise records are included. Species are sorted from most to least number of precise records (same order as in Fig. 3). Note the log scale.

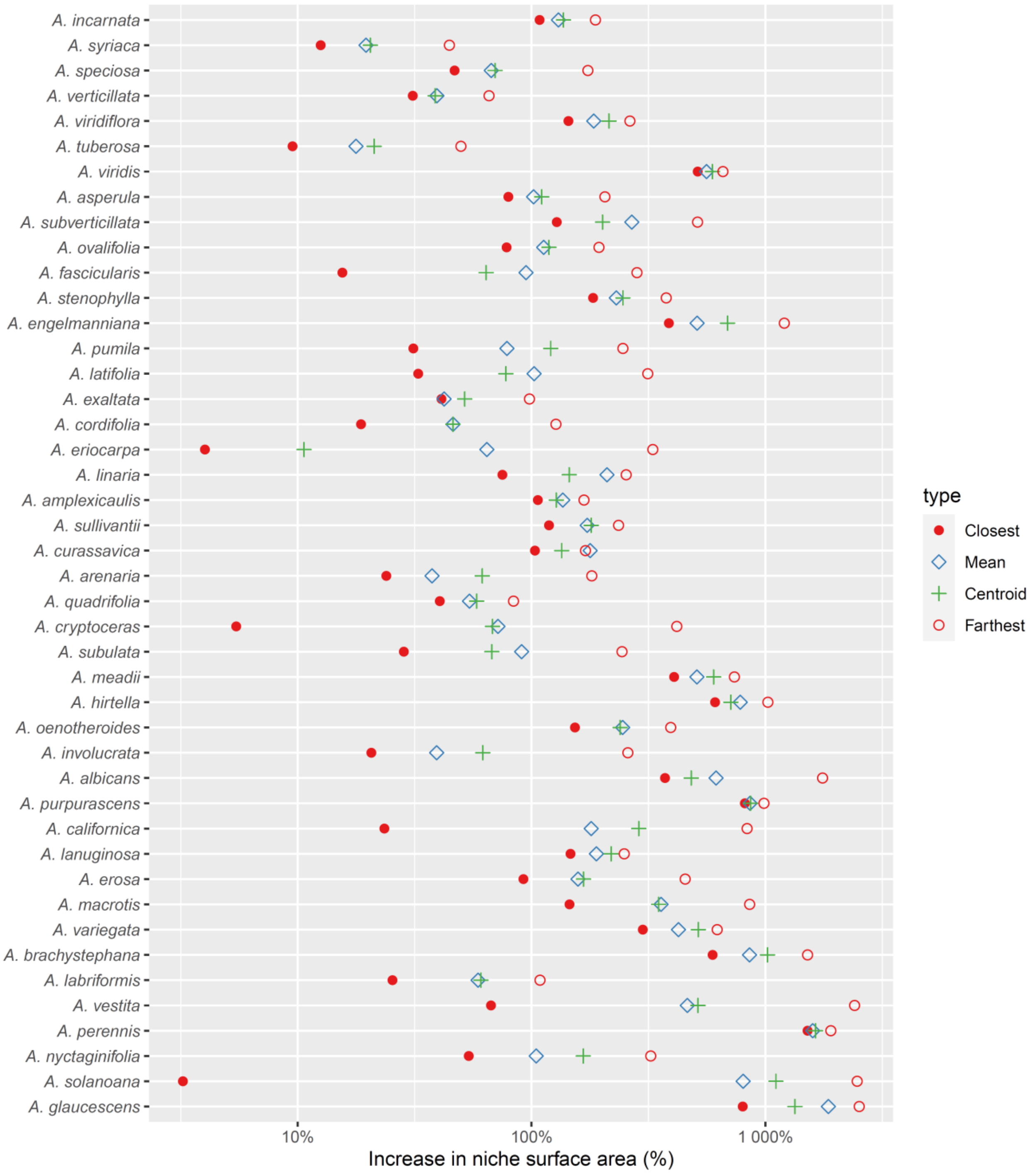


**Figure S3.4** Increase in **multivariate niche surface area** when imprecise records are included. Species are sorted from most to least number of precise records (same order as in Fig. 3). Note the log scale.

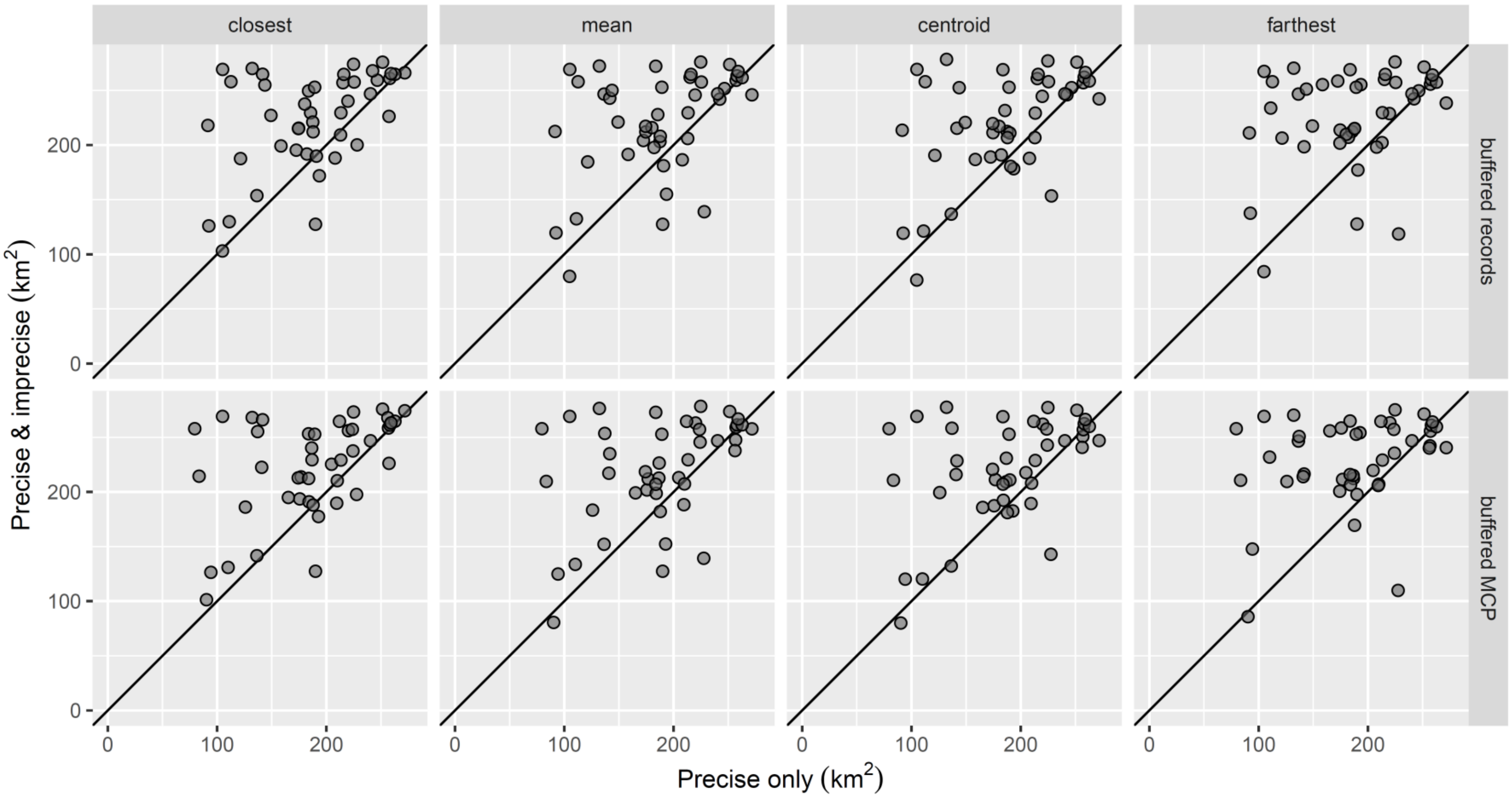


**Figure S3.5** Change in **present-day climatically suitable area** when imprecise records are included using Maxent models calibrated on a training region encompassing either 300-km buffers around each record or around the minimum convex polygon (MCP) encompassing all records. Species are sorted from most to least number of precise records (same order as in Fig. 3).

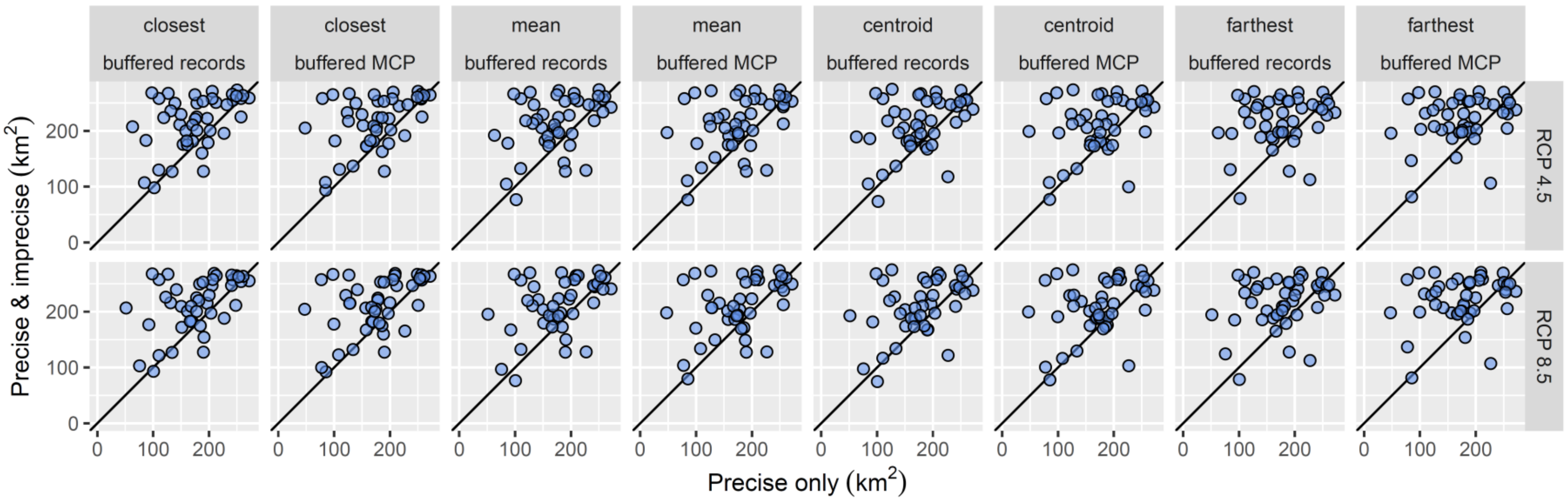


**Figure S3.6** Difference in **stable climatically suitable area** between ecological niche models using only precise records versus precise plus imprecise records. Panels display results based on Maxent models calibrated on a training region encompassing either 300-km buffers around each record or around the minimum convex polygon (MCP) encompassing all records and projected to RCPs 4.5 and 8.5 in the 2070s.

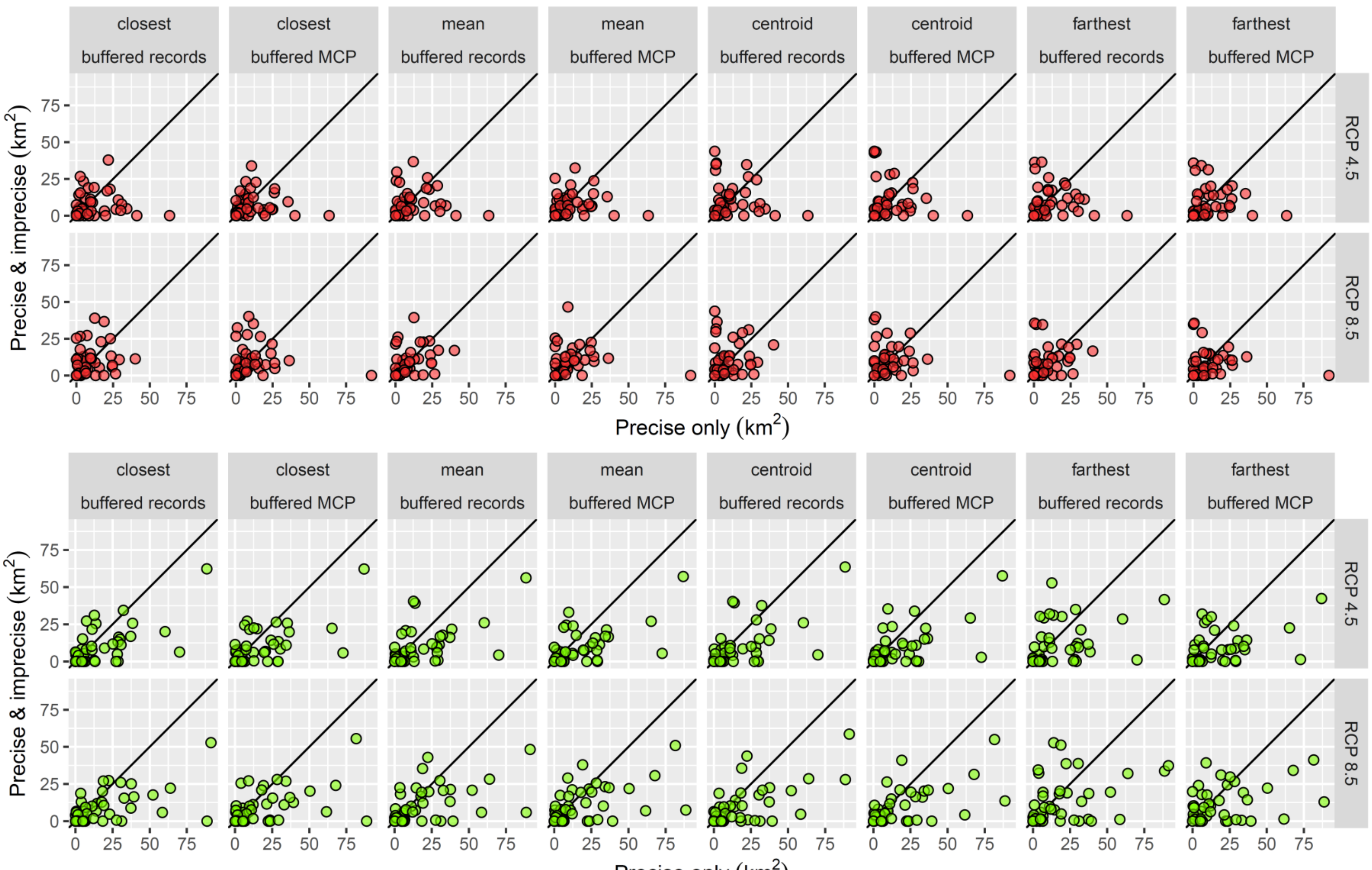


**Figure S3.7** Difference in **lost (top) and gained (bottom) climatically suitable area** between ecological niche models using only precise records versus precise plus imprecise records.
